# ETV2 mediated differentiation of human pluripotent stem cells results in functional endothelial cells for engineering advanced vascularized microphysiological models

**DOI:** 10.1101/2025.10.01.679558

**Authors:** Shun Zhang, Zhengpeng Wan, Lei Wang, Caihong Wu, Junkai Zhang, Sarah Spitz, Xun Wang, Marie A. Floryan, Mark F Coughlin, Francesca M. Pramotton, Liling Xu, Ron Weiss, Roger D. Kamm

## Abstract

Patient-specific microphysiological models, exemplified by organs-on-a-chip and organoids, have become a valuable tool for broad applications, revolutionizing biomedical research. However, limitations persist, with functional vasculature being a significant challenge. Generating functional human induced pluripotent stem cell (h-iPSC) derived endothelial cells (h-iECs) represent an urgent need. With the discovery of ETV2’s determinant role in specifying EC lineages during differentiation, researchers have adopted techniques involving ETV2 overexpression to produce h-iECs more efficiently and consistently. However, the capacity of these cells to form functional vasculatures has not yet been thoroughly investigated. Here, we generated multiple h-iPSC lines with inducible ETV2 expression, and subsequently differentiated them into h-iECs, which were validated functionally and by key endothelial markers and RNA-seq analysis. These cells are capable of self-organizing into stable microvascular networks (MVNs) in a microfluidic chip reproducibly, forming lumenized and functional vessels that mimic the in vivo capillary bed in both morphology and function – a result not achieved using h-iECs differentiated with conventional two-step methods using the same h-iPSC lines. Furthermore, complex microphysiological models featuring perfusable vasculature were also successfully developed using ETV2 activated h-iECs, demonstrated with vascularized tumor and blood-brain barrier (BBB) models. Additionally, by pooling genetically engineered h-iPSCs with inducible ETV2, we effectively employed an orthogonally induced differentiation approach to enhance vascularization of an organoid model. Our methodology opens avenues in precision medicine, leading to personalized microphysiological models with perfusable vasculature for various applications.

## 1. Introduction

Recent decades have witnessed an explosive growth in the development and application of in vitro microphysiological models, revolutionizing the field of biomedical research. Two prominent examples are organ-on-a-chip and organoid models (1, 2). Despite fundamental differences in design and engineering principles, both approaches aim to build multicellular in vitro organotypic models recapitulating the physiology and functionality of corresponding organs. They have already demonstrated tremendous potential in drug development, regenerative therapy, personalized medicine and mimicking organ level development and disease, bridging the gap between traditional cell culture models and whole-animal experiments (3–6). Despite these considerable achievements and rapid evolution, this field still faces limitations and unmet needs that hinder its wider application and limit broader impact. One major challenge is the lack a vasculature with consistent form and function (7–9). For example, without a perfusable vasculature, nutrients and oxygen cannot efficiently reach the innermost regions of many organoid models, leading to the formation of a necrotic core, which hinders proper cellular function and compromises the overall viability and functionality of the model. Additionally, since vasculature is involved in almost all physiological and pathological events, it is indispensable for mimicking higher-level organ functions such as the innate or adaptive immune response mediated by circulating cells and facilitating multi-organ interactions.

Endothelial cells (ECs), which form the interior lining of blood vessels, are an essential component of vasculature. Advances in human induced pluripotent stem cells (h-iPSCs) have opened exciting avenues to generate patient-specific ECs for engineering vascularized autologous microphysiological models (10). Conventional differentiation methods, also known as directed differentiation, involve a two-step process. Initially, h-iPSCs are differentiated into mesodermal progenitor cells (hMPCs) by activating Wnt pathways, followed by a vascular specification step utilizing vascular endothelial growth factor (VEGF) signaling (11, 12). However, those protocols suffer from certain limitations. Firstly, the yield of differentiated ECs is generally low, with most reported protocols achieving less than a 20% yield of ECs in the differentiated cell populations (12, 13). Additionally, achieving consistent and reliable differentiation across different h-iPSC lines remains challenging, likely due to the biological variabilities inherent in each h-iPSC line and the use of undefined components, such as fetal bovine serum (FBS), during culture or differentiation (14). ETS variant transcription factor 2 (ETV2) has been reported as a crucial regulator of vascular cell development (15–18). Following the pioneering work of Morita et al, who demonstrated the trans-differentiation of human fibroblasts into functional endothelial cells through ETV2 activation, extensive research has focused on understanding the role of ETV2 in specifying cells towards endothelial and hematopoietic lineages (19–23). ETV2 activation has also been combined with orthogonal forced overexpression of other transcription factors (TFs) to co-differentiate h-iPSCs into distinct cell types simultaneously, facilitating the engineering of vascularized tissues or organoids (24). Several h-iEC differentiation protocols involving ETV2 overexpression have also been proposed, either through temporal delivery of modified mRNA encoding ETV2 or by introducing a doxycycline-induced ETV2 expression system (25, 26). Both approaches achieved exceedingly high efficiency (>90%) in differentiating various human iPSC lines to ECs (25, 26). Previous studies have demonstrated that that ETV2 overexpression enhances the vasculogenic capacity of ECs (27), and the functional competence in forming functional MVNs in vitro. However, the enormous potential of transient expression protocols with a variety of iPS cell lines and in a variety of settings is still under investigation and represents a continuing need to establish the full range of applications.

In this work, we adapted previous protocols (23, 25, 26) to establish multiple h-iPSC lines with inducible ETV2 and subsequently differentiated them into h-iECs in a robust and efficient manner. We demonstrate the functional competence of these h-iECs by successfully generating capillary-like perfusable MVNs in vitro with several commonly used iPS cell lines, which cannot be formed using h-iECs derived from the same lines using the conventional two-step differentiation method. Moreover, we further establish the broad applicability of those functional h-iECs by i) developing a vascularized tumor model to evaluate chimeric antigen receptor (CAR) T-cell killing efficiency, and ii) demonstrating enhanced internal vascularization of a liver organoid using an orthogonally induced differentiation approach (28).

## 2. Results

### 2.1. Efficient and robust differentiation of h-iECs through transient activation of ETV2

Building on previous studies (23, 25), we have adopted a differentiation protocol using doxycycline (Dox)-induced ETV2 overexpression in pre-programmed h-iPSCs. While a recent study by Luo et al. (26) reported a similar approach, our work differs in ETV2 constructs, transfection method, and replating strategies. Two types of plasmids were constructed and transfected with corresponding methods into various h-iPSC lines (Figure S1a). Consistent with the methods developed previously, we first differentiated h-iPSCs into hMPCs through a two-day treatment of the glycogen synthase kinase 3 inhibitor CHIR99021 (Figure 1a,f), followed immediately by a 24-hour induction of ETV2 overexpression via Dox, in combination with VEGF, FGF, and other growth factors to promote the conversion of hMPCs into h-iECs (Figure 1a).

**Figure 1.**
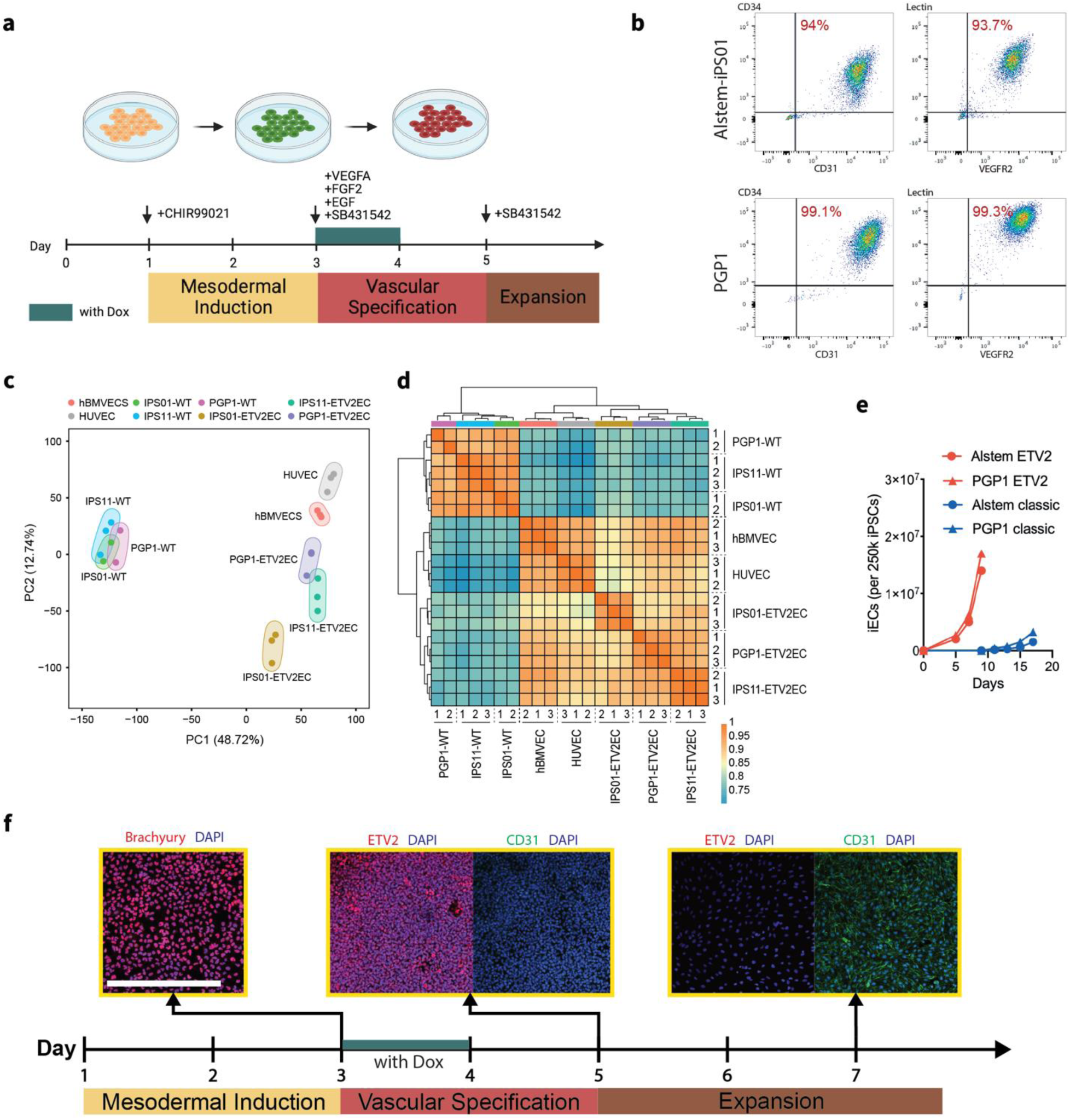
Efficient and robust differentiation of h-iECs through transient activation of ETV2. (a) Schematic of two step h-iECs differentiation protocol with transient activation of ETV2. Details can be found in methods section. (b) Differentiation efficiency of h-iPSCs into CD31^+^/CD34^+^/VEGFR2^+^/ UEA-I ^+^ h-iECs with ETV2 transient activation by flow cytometry. (c) Principal component analysis and (d) hierarchical clustering with pairwise correlation for bulk RNA-seq analysis on various iECs, primary ECs and hiPSCs (n=2 for PGP1 and iPS01 iPSC lines, n=3 for all other groups). (e) Expansion curve of h-iECs using two step method with or without ETV2 activation, all started with 250k h-iPSCs. (f) Immunostaining of characteristic markers during EC differentiation using two step method with ETV2 activation. Scale bar is 500 μm.

We first compared our transient protocol with the conventional 10-day differentiation approach (12), which usually yields differentiation efficiencies of less than 20% (Figure S1b). Consistent with Luo et al (26) we found that transient activation of ETV2 significantly enhanced the efficiency of differentiation. We used multiple EC markers, including CD31, CD34, VEGFR2 and UEA-I, to characterize h-iECs. By day 7, a majority of the cells had successfully differentiated into EC lineages (Figure 1b). Notably, the yield of h-iECs was substantially improved, resulting in a more than tenfold increase in the number of ECs compared to the conventional, two-step method without transient ETV2 activation (Figure 1e). The addition of Dox to the culture media facilitated rapid and uniform expression of ETV2 (Figure 1f), which is crucial for efficient differentiation. Immunostaining of those h-iECs at day 7 during differentiation revealed abundant and uniform expression of the EC marker CD31 (Figure 1f), consistent with flow cytometry data.

To identify potential differences between h-iECs generated using our protocol and primary ECs, we conducted bulk RNA-seq analysis on all three h-iEC samples differentiated through transient activation of ETV2. The undifferentiated parental h-iPSCs were included as negative controls, while human umbilical vein endothelial cells (HUVECs) and human brain microvascular endothelial cells (hBMVECs), two primary EC types utilized in this study to form MVNs, served as positive controls. Principal component analysis (PCA) revealed that various h-iECs clustered more closely with primary HUVECs and hBMVECs, comparing with their parental h-iPSCs, indicating a successful transcriptional transition towards an endothelial phenotype (Figure 1c), which is further substantiated by hierarchical clustering and pairwise correlations (Figure 1d).

### 2.2. Transient activation of ETV2 renders h-iECs with enhanced vasculogenesis capacity to form functional MVNs in vitro

We next assessed h-iECs’ functionality, specifically focusing on their capacity to form capillary-like vascular networks in vitro when co-cultured with supporting cells (human lung fibroblasts, HLFs, in this case) (Figure 2a). None of the h-iECs differentiated with the conventional two-step method without ETV2 activation formed well-connected vascular networks, and no patent lumens could be identified with dextran perfusion (Figure 2b, Figure S2a). In contrast, h-iECs differentiated with transient ETV2 activation self-organized into stable lumenized MVNs, as evidenced by successful dextran perfusion (Figure 2c, Figure S2). It is worth mentioning that due to the remarkably high differentiation efficiency resulting from ETV2 activation, we were able to utilize the differentiated cells directly to form MVNs without the need for cell sorting. Additionally, we found that h-iEC MVNs formed with high serum (10% FBS) culture medium were more robust and fully perfusable than ones formed with 2% FBS (Figure 2c). Characteristic markers of ECs and ECM (CD31, VE-CAD, ZO-1 and collagen IV) are clearly evident in the vascular networks formed within the device (Figure 2d). When we characterize the morphological parameters of MVNs, we found that the networks formed with h-iECs exhibited smaller vessel diameter with more branches compared to those formed with HUVECs, the most commonly used ECs for in vitro MVNs models (Figure 2e-h). These morphological metrics of narrower lumens and greater branching density are hallmarks of capillary architecture, suggesting the formation of more capillary-like vascular bed with h-iECs.

**Figure 2.**
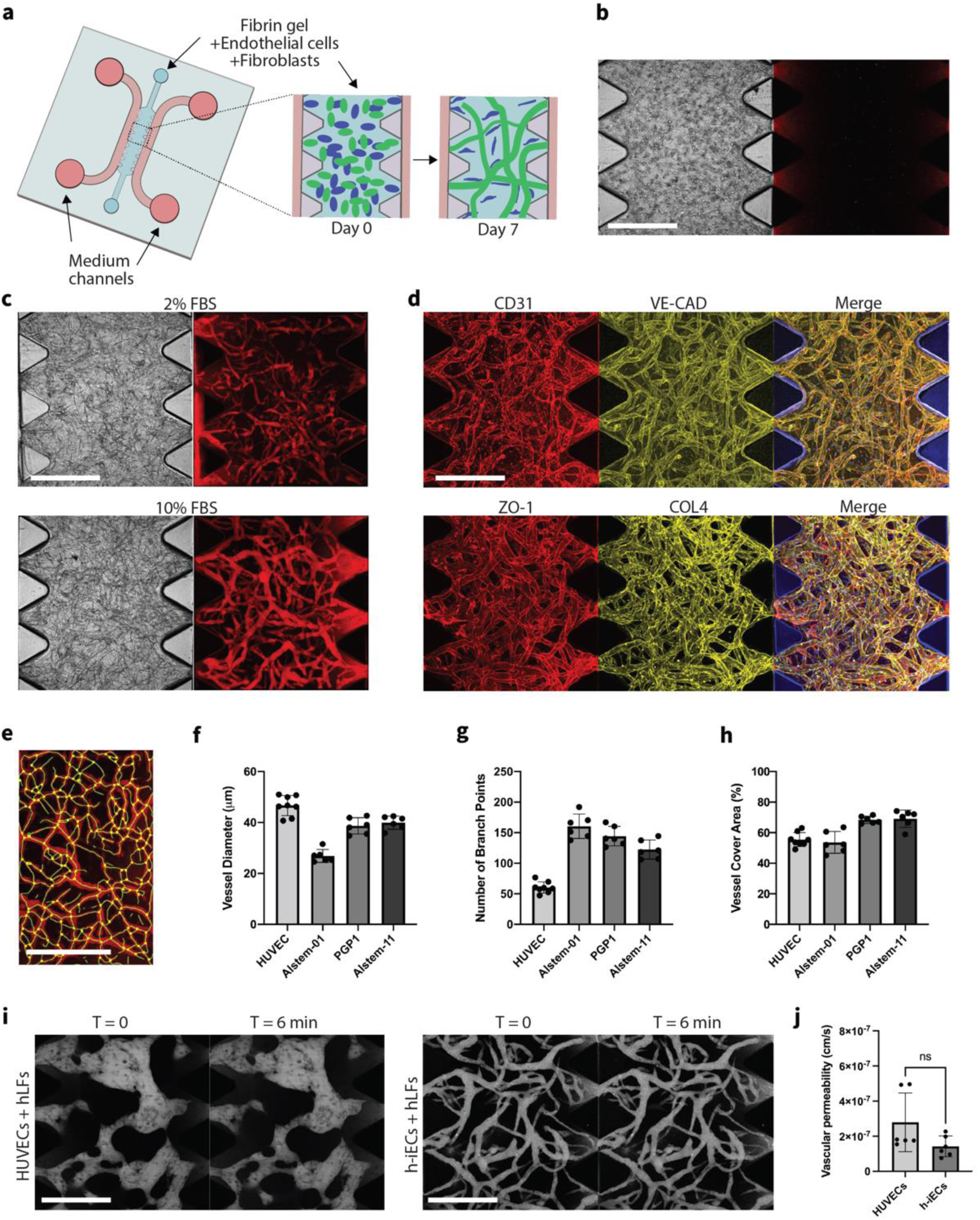
Formation of perfusable MVNs with h-iECs differentiated through transient activation of ETV2. (a) Schematic diagram of the AIM Biotech chip used to generate MVNs with h-iECs and HLFs mixtures encapsulated in fibrin gel. (b) Representative images of MVNs made of h-iECs on day 7. h-iECs were differentiated from Alstem iPS01 h-iPSCs with conventional methods. Perfusion test was performed with 40 kDa Texas Red dextran (red). (c) Representative images of MVNs made of h-iECs on day 7, supplemented with 2% or 10% FBS. h-iECs were differentiated from Alstem iPS01 h-iPSCs engineered with inducible ETV2 using our optimized protocol. Perfusion test was performed with 40 kDa Texas Red dextran (red). (d) Immunofluorescence staining for various proteins of interest in the perfusable MVNs with h-iECs differentiated with our optimal protocol, including CD31, VE-cadherin, ZO-1 and Collagen IV. (e) Perfusable vessel skeleton and junction points detected using fluorescent image acquired for dextran. (f-h) Statistical analysis of morphological parameters of perfusable MVNs formed with various EC sources. n = 8 devices for HUVEC, n = 6 devices for each h-iECs differentiated with our optimal protocol. (i) Vascular permeability measurements for perfusable MVNs engineered using HUVECs or h-iECs, with hLFs. Confocal z-stack images of perfusable MVNs with dextran were acquired with 6 min time interval. Collapsed z-stack image are shown. (j) Vascular permeability measurements of 40 kDa dextran for perfusable MVNs. n=2 devices for each group, 3 measurements were performed in each device at different ROIs. All scale bars are 500 μm.

To further evaluate the functionality of the engineered MVNs, we measured vascular permeability, a critical aspect of their functionality, by perfusing 40 kDa dextran in MVNs formed by HUVECs or h-iECs, both with hLFs as supporting stromal cells. Fluorescence imaging at 0 and 6 min post-perfusion revealed only a slight increase in background fluorescence, indicating good permeability within the networks (Figure 2i). We found h-iEC-derived MVNs exhibit similar permeability to HUVEC-based MVNs (Figure 2j), highlighting the functional capability of the differentiated h-iECs.

Next, we further explored the application of these functional h-iECs by examining their potential for forming other organotypic MVN models. Following the protocol that we developed previously (29), we engineered an in vitro BBB model consisting of h-iECs, co-cultured with human primary brain pericytes (PCs), and astrocytes (ACs). h-iECs developed into a fully perfusable MVN with PCs and ACs residing in the interstitial space surrounding the microvessels (Figure S2d). PCs and ACs were in direct contact with endothelium, as described in other similar brain MVN models (30, 31).

### 2.4 Perfusion of h-iECs with immune cells

We next tested the vasculogenesis capacity of these h-iECs differentiated with transient ETV2 activation seeded in a much larger microfluidic device with a circular (5mm diameter) central region. Well-connected vascular networks still formed across the entire gel region, as confirmed by the successful perfusion of peripheral blood mononuclear cells (PBMCs) (Figure 3a). Functionality of the MVNs was further confirmed with the extravasation of PBMCs into the extracellular space through vessel walls. Moreover, PBMCs pre-treated with phytohemagglutinin (PHA) lead to significantly increased extravasation events, which suggests the PBMC-EC interaction can be faithfully recapitulated with this system (Figure 3b-d).

**Figure 3.**
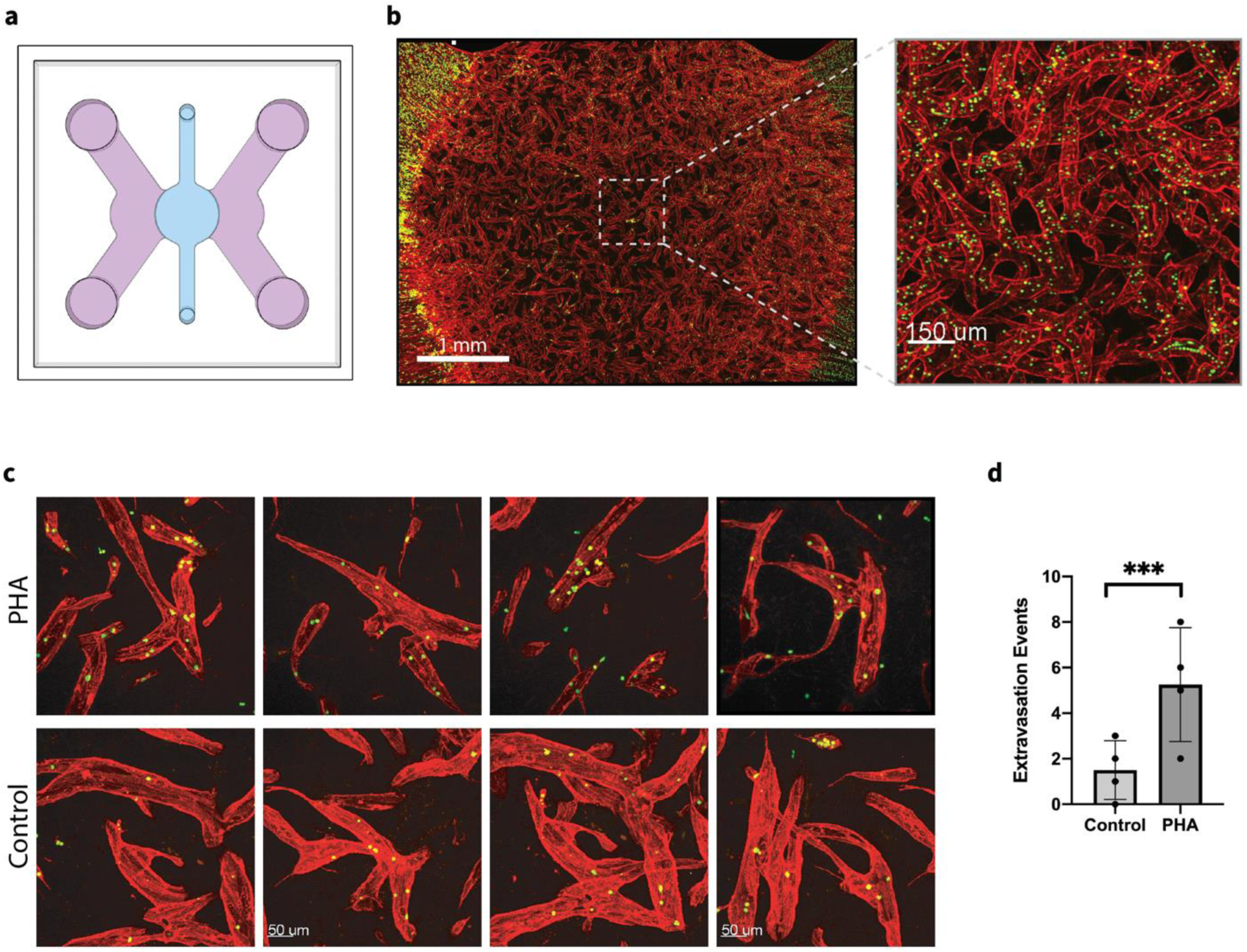
Formation of bulk MVNs with PBMCs perfusion. (a) Schematic diagram of the microfluidic chip used to generate MVNs with HLFs and h-iECs differentiated from Alstem iPS11 h-iPSCs engineered with inducible ETV2 using our optimized protocol. (b) Representative images of MVNs formed in customized microfluidic chip at day 7 and the perfusion of PBMCs, with zoomed in view. Red: UEA-I live staining of h-iECs, Green: PBMCs. Scale bar is 1 mm for the left panel, and 150 µm for expanded view. (c) High magnification images (with 30x objectives) of MVNs after perfusion of control PBMCs and PBMCs pretreated with PHA. Scale bar is 50 µm. (d) Statistical analysis of extravasation events of control PBMCs and PBMCs pretreated with PHA in MVNs. n = 4 devices for each case. Significance was calculated with t-tests. ***P < 0.01.

### 2.4 Vascularized tumor spheroid model with h-iECs

Many groups, including our own, are currently developing methods to grow perfusable tumoroids or tumor cell spheroids for studies of cancer metastasis (32–36). We therefore sought to demonstrate a vascularized tumor-on-a-chip model to investigate the anti-tumor activity of CAR-T cells. To facilitate the formation of open vessels between micro-posts and capillary-like MVNs throughout the central gel region with robust development of fully perfusable MVNs, we adopted a previously reported 2-step seeding strategy (as illustrated in Figure 4a) (37). We observed the formation of a dense capillary-like vessel network with connected patent lumens surrounding the embedded tumor spheroid, as evidenced by successful dextran perfusion (Figure 4b). Although the majority of the vessels were at the periphery of the tumor spheroid, we also observed vessels sprouting into the tumor region, effectively recapitulating certain aspects of the tumor microenvironment in vitro.

**Figure 4.**
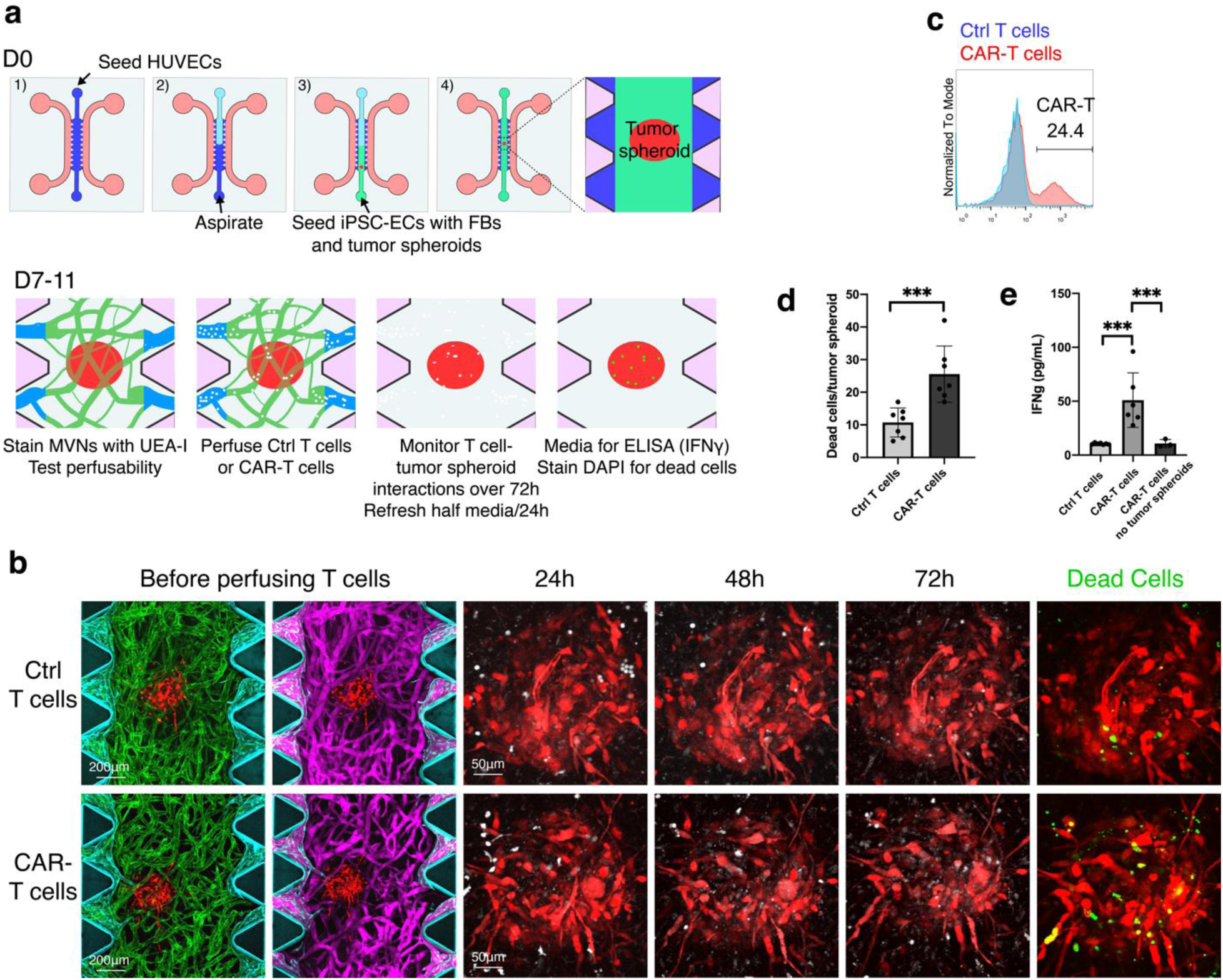
CAR-T killing assay of vascularized tumor model. (a) Schematic illustration of CAR-T cell killing assay in vascularized tumor spheroid model made with 2-step seeding strategy. HUVECs were used as outer layer ECs to increase the chance of MVNs openings forming between pillars. h-iECs, HLFs, and tumor spheroid were co-seeded to establish central vascularized tumor spheroid structure. On day 7, after MVNs containing tumor spheroid were stained with UEA-I and perfused with dextran to detect morphology and perfusability, control T cells or CAR-T cells were perfused to MVNs to examine the T cell-tumor cell interaction and killing. On day 11, conditioned media were collected for cytokine test (interferon gamma, IFNγ), and then devices were stained with DAPI to visualize dead cells. (b) Representative confocal images of vascularized Skov3 tumor spheroids treated with control T cells or mesothelin specific CAR-T cells. Before perfusing T cells: cyan, HUVECs (outer layer); total ECs were stained with UEA-I (green); magenta, dextran; red, Skov3 tumor spheroid. After perfusing T cells: white, T cells; red, Skov3 tumor cells; green, DAPI staining in live sample to illustrate dead cells. Scale bar is 200 µm for left panels and 50 µm for expanded view of tumor spheroids. (c) Histogram showing expression of anti-mouse F(ab)2 to demonstrate percentage of anti-mesothelin specific CAR-T cells. (d) Statistical analysis of dead cells in vascularized tumor spheroids treated with control T cells or CAR-T cells. n = 7 devices for each case. Significance determined by t-tests. ***P < 0.01. (e) IFNγ concentrations of conditioned media from vascularized tumor spheroids treated with control cells (n = 7 devices), CAR-T cells (n = 7 devices), or from MVNs alone perfused with CAR-T cells (n = 3 devices). Significance determined by t-tests. ***P < 0.01.

Control T cells or CAR-T cells (with an approximate 25% CAR positive rate, Figure 4c) were perfused into the media channel, where they then entered into the tumoral MVNs, extravasated, and interacted with tumor cells. We assessed the T cell response using dead cell staining and cytokine secretion analysis. 72 hours after perfusion, we observed higher dead cell densities in the CAR-T cell groups compared to the control T cell group (Figure 4d). Media from devices were analyzed for IFNγ, an indicator of immune activation and antitumor response, using the enzyme-linked immunoassay (ELISA). The results indicated a stronger response from CAR-T cells in the MVNs with an embedded tumor spheroid, compared to control T cells or CAR-T cells in MVNs lacking tumor (Figure 4d). Collectively, these data demonstrate the efficacy of this vascularized tumor spheroid model in evaluating CAR-T cell responses to tumor cells in vitro. It serves as a valuable platform for studying the interactions between immune cells and tumors in a more physiologically relevant microenvironment and also offers the prospect of creating an isogenic model if patient-derived iPSCs and immune cells are used.

### 2.5 Enhanced vascularization of a liver organoid model through orthogonal differentiation

ETV2-expressing pluripotent stem cells have been utilized to vascularize various organoid models (38). Here, we aim to develop complex vascularized organoid models by simultaneously co-differentiating h-iPSCs through overexpression of certain TFs. Previously, Guy et al reported a 2-D method to generate a liver bud-like structure containing hepatocytes, ECs and stromal cells by inducing a wide range of GATA6 expression in a pluripotent cell population using a Dox-inducible system (28). We first sought to reproduce a similar multicellular liver bud-like structure in a 3D cell aggregate by inducing heterogeneous expression of GATA6 (Figure 5a). Notably, our results showed a clear expression of CEBPα, CD31 and desmin (Figure 5c, Figure S3), indicating the successful formation of a liver bud-like niche consisting of hepatocytes, ECs and stellate cells. However, we observed that ECs only formed in small, scattered clusters and failed to establish well-connected vascular networks.

**Figure 5.**
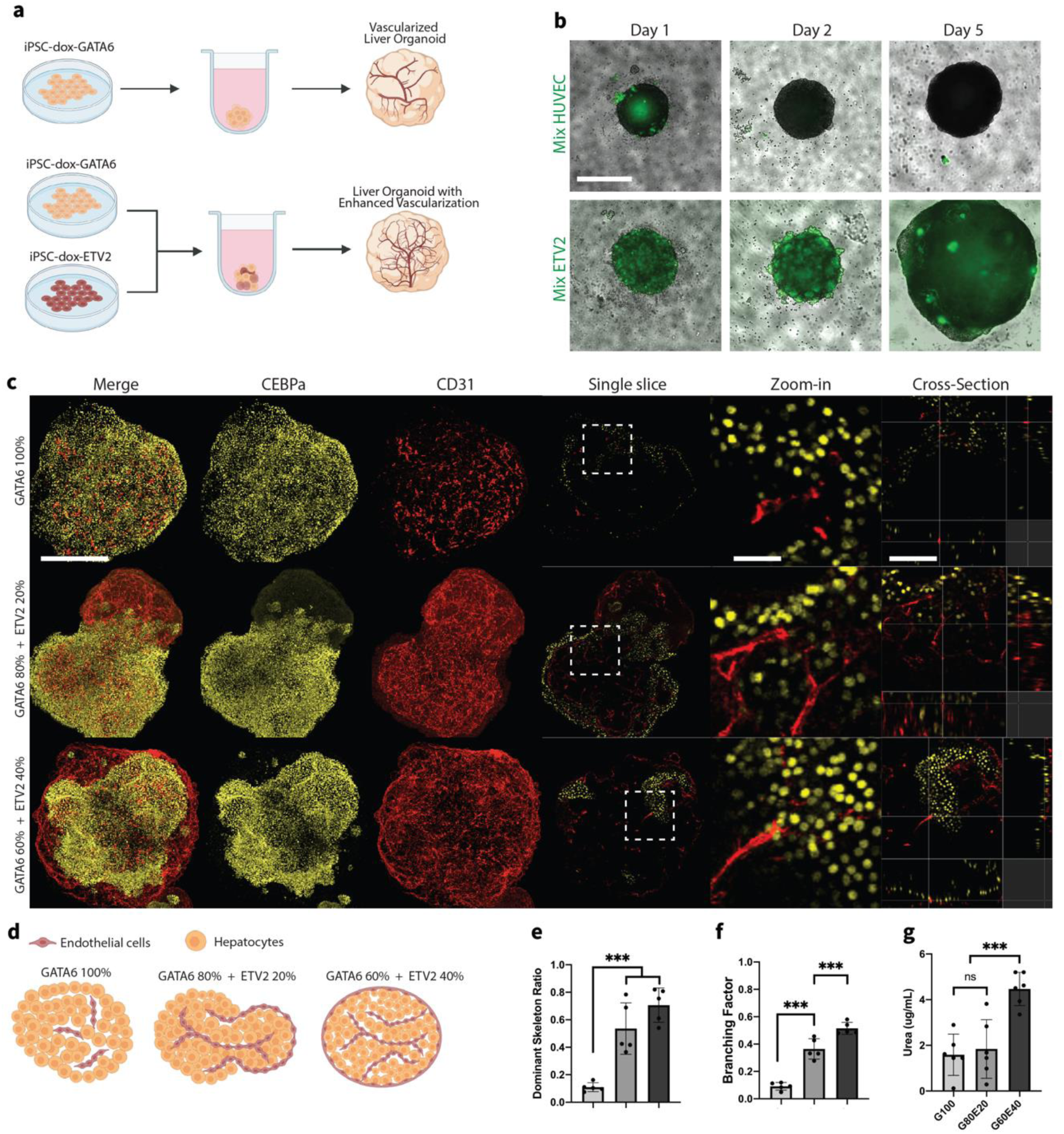
Generation of liver organoid model with enhanced vascularization. (a) Schematic view of the methods for generating liver organoids. (b) Embryoid body formation. 1 μg ml^−1^ doxycycline was added from day 0. Top: 80% PGP1-GATA6 and 20% GFP-HUVEC pooled embryoid body. Green, GFP HUVECs. Bottom: 80% PGP1-GATA6 and 20% PGP1-ETV2 pooled embryoid body. Green, induced PGP1-ETV2-EYFP. Scale bar is 500 μm. (c) Fluorescent immunostaining images of CEBPa (hepatocyte marker) and CD31 (endothelial cell marker) in day 22 liver organoids generated by pooling varying ratios of PGP1-GATA6 and PGP1-ETV2 cells. The first three columns show confocal z-stack projections, followed by representative single-slice images and corresponding zoom-in views. Cross-sectional views are displayed in the rightmost column. Scale bars: 500 μm (z-stack projections and single slices), 100 μm (zoom-in views), and 200 μm (cross-sectional views). (d) Schematic illustration of spatial arrangements of ECs and hepatocytes within liver organoid by day 22. Organoids were formed by pooling different ratio of PGP1-GATA6 and PGP1-ETV2 initially. (e,f) Statistical analysis of morphological parameters of vascular network within day 22 liver organoids. n = 5 organoids for each case. Significance was calculated with t-tests. ***P < 0.01. (g) Statistical analysis of urea production from day 22 liver organoids. n = 6 organoids for each case. Significance was calculated with t-tests. ***P < 0.01.

To enhance vascularization and improve vascular network connectivity within the organoids, we mixed GATA6-h-iPSCs with HUVECs, which are a commonly used endothelial cell type for vascularizing organoid models (39–41). However, this led to a complete separation of the two cell types within 2 days (Figure 5b), possibly due to differences in cell adhesion molecules. Next, we adopted a new approach by pre-mixing GATA6-h-iPSCs with ETV2-h-iPSCs in one-pot culture conditions, similar to the orthogonal differentiation method established by Skylar-Scott et al (24). When GATA6 and ETV2 h-iPSCs were mixed and co-cultured, single integrated embryoid bodies were formed with interspersed cells (Figure 5b).

We then produced liver organoids with improved vascularization by pooling isogenic GATA6-h-iPSCs and 20% or 40% ETV2-h-iPSCs to form embryoid bodies with a 5-day transient activation of pre-programmed TFs. At day 22, compared with GATA6-only organoids, organoids composed of mixed cell populations exhibited extensively enriched vascular networks, as marked by CD31 (Figure 5c, Figure S4). Zoom-in and cross-sectional images further revealed that in GATA6-only organoids, CD31⁺ signals were sparse and discontinuous, lacking organized structure. In contrast, organoids containing 20% or 40% ETV2-h-iPSCs developed continuous and branching CD31⁺ networks that resembled vascular structures. Notably, in these mixed organoids, we also observed the formation of lumen-like structures, indicative of endothelial tube morphogenesis and improved vascular maturation. Detailed characterization revealed that organoids composed of mixed cell populations had significantly improved vascular connectivity, as quantified by dominant skeletal ratio (proportion of the longest skeleton path relative to the total path length of all skeletons) and branching factors (the ratio between junction numbers and endpoint numbers), while maintaining the essential function of urea production (Figure 5e,f,g).

Interestingly, we found the initial cell composition directly affected the spatial distribution of the cells in the organoid, particularly affecting arrangement of hepatocytes and ECs. By day 22, all ECs in the GATA6-only organoids were found in the interior of the organoid, surrounded by hepatocytes that were distributed both at the periphery and throughout the organoid, as depicted in the cartoon and illustrated with single-slice immunostaining images (Figure 5c). However, when mixing with 20% ETV2 h-iPSC initially, the organoid self-organized into a dumbbell-like structure, with one side maintaining a similar polarity to that of the GATA6-only organoids. On the other side, the polarity was inverted, with ECs encapsulating hepatocytes. Increasing the initial ETV2 population to 40% resulted in the day 22 organoid adopting a spherical shape like GATA6-only organoids, but with a complete inversion of polarity. In this case, hepatocytes were located in the interior of the organoid, encapsulated by ECs. These distinct morphological patterns and cell distribution arrangements were consistently observed in all the experiments, supporting the trends we described here (Figure 5c, Figure S4).

## 3. Discussion

h-iECs hold great promise for a myriad of scientific and clinical applications in regenerative medicine and vascular biology. By offering a renewable and autologous source of ECs, h-iECs open new avenues for disease modeling, drug discovery, and personalized medicine, particularly when combined with advanced microphysiological systems (42, 43). Despite tremendous efforts devoted to the development of improved differentiation methods with higher efficiency and robustness, there remains a need for competent ECs to engineer microphysiological systems with functional MVNs (11, 44, 45). We demonstrate that the transient activation of ETV2 resulted in h-iECs with enhanced vasculogenesis capacity to form functional MVNs in a robust manner. The underlying mechanisms that underpin this enhanced vasculogenic capacity in h-iECs to form functional lumens remain elusive, since formation of MVNs is a highly orchestrated process involving multiple interrelated events, such as EC migration and proliferation, ECM remodeling, lumen formation and stabilization, etc. (46–48). Notably, using mature human ECs, Palikuqi et al demonstrated that transient reactivation of ETV2 ‘resets’ the vasculogenic memory of mature ECs to an early embryonic stage, converting these ECs into adaptable, vasculogenic cells. Further investigation revealed that ETV2 induces tubulogenic pathways via chromatin remodeling and RAP1 activation, which promotes the formation of durable lumens (18, 27).

Recent studies from the Melero-Martin group have extensively discussed the advantages of Dox-inducible ETV2 endothelial cell differentiation, highlighting its efficiency, reproducibility, technical improvements, developmentally relevant induction, and comparisons with alternative approaches, including piggyBac and other transfection systems (26). Here, we focus on the application of these cells in microphysiological systems. Currently, vasculature is often integrated into complex multi-culture systems to mimic higher-level organ functions or facilitate multi-organ interactions (36, 38, 39, 49). In these complex in vitro systems, the microenvironment (the choice of hydrogel scaffold, composition of culture medium and growth factors, etc.) must be carefully designed to support the development and growth of all the cellular components. That is, h-iECs must be able to form functional MVNs under sub-optimal conditions, presenting a considerable challenge for their vasculogenesis capacity. Our findings demonstrate that the h-iECs generated with our protocol offer clear advantages for engineering complex microphysiological systems featuring self-organized MVNs, as the brain MVN system and vascularized tumor spheroid model demonstrated in this study. The capacity of h-iECs derived with transient ETV2 induction to self-assemble into perfusable MVNs enables the physiological vascularization of advanced microphysiological systems integrated into micro- and macro-fluidic devices. While other cellular components in the models presented in this study may come from primary cells or established cell lines, our efforts represent a crucial step towards fully patient-specific vascularized models.

HUVECs have frequently been used as EC sources to incorporate vasculature into organoid models due to their accessibility and ease of manipulation. However, their typical application involves co-culturing with organoids or other cell types once they have reached a mature state, often resulting in limited vascularization (41, 50). To address this limitation, one approach is to synchronize the vascular development process with that of the organoid. Our attempts to combine HUVECs with pluripotent stem cells led to a complete phase separation during embryoid body formation, a critical stage in the development of numerous organoid models. Using the dox-inducible system offers advantages in the orthogonal differentiation of multiple cell types. Leveraging our understanding of specific TFs such as GATA6 and ETV2, which respectively guide the differentiation of liver-bud-like structures and ECs, we effectively engineered liver organoids exhibiting improved vascularization. This was accomplished by overexpressing GATA6 or ETV2 in a subset of cells within developing organoids. The ‘orthogonal differentiation’ approach holds great potential for broad applications in tissue engineering, especially facilitating the generation of vascularized organoids and tissues (24).

While this orthogonal differentiation strategy succeeded in boosting vascularization in liver organoid model, we did not observe continuous patent lumens, such as those we demonstrated with suspended cells in microfluidic devices. Consequently, the critical functionality of capillaries to provide sufficient nutrients and oxygen to nearby cellular components within the organoid model was not achieved. This shortcoming can be attributed, in part, to the fact that h-iECs differentiated in the organoid model did not adhere to the optimal differentiation protocols we have established, which involves the transient and timely activation of ETV2 after converting h-iPSCs into h-MPCs. Instead, ETV2 was continuously overexpressed for 5 days following the formation of embryoid bodies, bypassing the intermediate mesodermal stage. Wang et al have shown that although direct induction of ETV2 expression in h-iPSCs could generate h-iECs rapidly and efficiently, those cells were associated with impaired functionality regarding cell migration, angiogenesis and expansion (37). One potential solution to address this limitation is to develop a platform capable of sensing cell state and conditionally activating expression of TFs that automatically orchestrate the multistep differentiation towards desired cell types. Leveraging the close relationship between expression of certain miRNA species and cell state, we successfully engineered miRNA sensors that could conditionally activate certain TF based on the activity of endogenous cell-state specific miRNAs (51). Another possible solution is to incorporate proper mechanical stimuli. For instance, the beneficial role of wall shear stresses induced by diverse flow types on angiogenesis, vasculogenesis and 3D capillary morphogenesis has been confirmed by various studies (52–55). Furthermore, Homan et al. reported an in vitro method for culturing kidney organoids placed on top of a customized hydrogel under flow, which induced the formation of EC networks within and sprouting from the organoids (56). Those methodologies could potentially work in a synergistic manner with integration of microfluidic techniques, which offer the advantage of precise control over the in vitro microenvironment, ultimately achieving the milestone of developing functional, perfusable vasculature within organoid models.

We found a close relationship between initial cell composition and spatial distribution of various cells in a mature liver organoid, shedding light on the complex cellular interactions that drive organoid development and intricately shape its spatial arrangement. Furthermore, this inversion of polarity holds the potential to establish full integration of pre-vascularized organoids with an in vitro capillary bed to achieve functional anastomoses, thereby enabling continuous perfusion. For GATA6-only organoids, all ECs resided in the interior of the organoid. In this scenario, both the internal vasculature of the organoid and the external vasculature of in vitro MVNs faced challenges in breaching the ‘shell’ of the organoid to establish functional connections. In contrast, when mixing with 40% ETV2-h-iPSC initially, a complete polarity inversion occurred, resulting in the outer ‘shell’ being composed of ECs. This configuration allows for direct physical contact between the vasculature of the organoid and the outer capillary bed, promoting the potential for spontaneous anastomosis.

## 4. Conclusion

We differentiated h-iECs through transient activation of ETV2, which consistently yields remarkably high differentiation efficiency among all the tested h-iPSC lines. Furthermore, we assessed the functional competence of these h-iECs by successfully generating capillary-like, perfusable MVNs in vitro, which cannot be obtained through conventional h-iEC differentiation methods using the same h-iPSC lines. To demonstrate the application of those competent h-iECs in engineering complex vascularized microphysiological systems, we developed a vascularized tumor model for assessing CAR-T cell killing efficiency, as well as a BBB model. Additionally, by pooling genetically engineered h-iPSCs with inducible TFs, we effectively employed an orthogonally induced differentiation approach to develop liver organoid models with enhanced vascularization. These methods could have broad applications in precision medicine, enabling the development of autologous vascularized microphysiological models for comprehensive drug and treatment evaluations. Furthermore, its potential extends to producing clinical-grade h-iECs for various regenerative therapies.

## 5. Experimental Section/Methods

### TF-induced h-iPSC cell lines

#### Plasmids and transfection

Human ETV2 was cloned into the pCW57-GFP-P2A-MCS (Neo) plasmid using Gibson assembly method to construct pCW57-GFP-P2A-ETV2 plasmid. ETV2 was cloned from pSIN4-EF1a-ETV2-IRES-Puro plasmid, a gift from Igor Slukvin (Addgene plasmid # 61061; http://n2t.net/addgene:61061; RRID:Addgene_61061) (57). pCW57-GFP-P2A-MCS (Neo) was a gift from Adam Karpf (Addgene plasmid # 89181; http://n2t.net/addgene:89181; RRID:Addgene_89181) (58). The pCW57-GFP-P2A-ETV2 construct was co-transfected with psPAX2, pMD2.G for packaging into HEK293T cells and the supernatant of the HEK293T culture was collected on day 2 and 3 post-transfection, followed by Lenti-X™ Concentrator (Takara Bio) mediated concentration. Alstem iPS01 or iPS11 cells were incubated with the concentrated lentivirus in the presence of polybrene, 8 µg/ml, overnight. Cells were then selected with G418 (Invivogen) for a week. Transfected iPSCs were dissociated into single cells and subsequently sorted using BD FACSAria™ III Cell Sorter to form single clones in 96-well plate. Single clones were duplicated and assessed for successful transfection by detecting GFP signals induced by addition of 1 μg/mL doxycycline for 24 hours. Successfully transfected iPSC clones were expanded and used for subsequent experiments.

The modular PiggyBac plasmid cloning scheme used in this study is based on the extended MoClo scheme published previously (59, 60). In brief, genetic elements are first cloned into Level 0 plasmid backbones corresponding to Insulators (pL0-I), Promoters (pL0-P), 5’ UTR (pL0-5), Gene (pL0-G), 3’ UTR (pL0-3) and Terminators (pL0-T). Then the genetic elements can be cloned into positioning backbones for transcriptional unites (pL1s), with ST1-2, ST2-3 and ST3-4X for position 1, 2 and 3, respectively. To make a genetic circuit to express ETV2 in an inducible manner, the ST1-2 backbone contains a TRE3 inducible promoter driving the expression of ETV2-P2A-EYFP, the ST2-3 backbone harbors a constitutive promoter for expressing rtTA3, and the ST3-4X backbone encodes a hygromycin resistance gene. Finally, the pL1s can be cloned into Level 2 vectors which includes a regular vector for transfection-based assays or PiggyBac transposon vector that can be stably integrated into the chromosome. The PiggyBac Inverted Terminal Repeats (ITRs) sequence was obtained as PCR products from AddGene (plasmid #40973) and were cloned into the flanking areas of the insertion site on the pL2 backbone using an infusion protocol (Takara Bio USA) (61). PiggyBac transposase was reconstructed based on the sequence published previously as a double-stranded gBlock and cloned into pL0-G (62). The sequences of transcriptional factor and proteins were based on UniProt.org or previous publication, the DNA sequences were ordered as double-stranded gBlocks from IDT and cloned into pL0-G.

Genetic circuits were stably integrated into PGP1 h-iPSCs by a PiggyBac system using Lipofectamine Stem (Invitrogen) in a 24-well format following manufacturer’s protocol. Complexes were prepared in Opti-MEM (Gibco). h-iPSCs were trypsinized, counted and plated on 24-well plate the day before transfection with the concentration of 4×10^4^ cells per well in 0.5 ml of complete growth medium. Cells should be 50∼80% confluent on the day of transfection. The cells are then incubated at 37°C and 5% CO_2_ for 18-24 hours post-transfection before assaying for transgene expression. A plasmid encoding the PiggyBac transposase was co-transfected with the transposon vectors in a molar ratio of 1:3, while maintaining a total DNA amount of 500 ng per well in a 24 well plate format. Two days after transfection, Hygromycin B (InvivoGen) was added to the mTeSR Plus medium at a concentration of 100 μg/mL. This selection step was conducted over a period of two weeks to enrich the population of h-iPSCs expressing the transgenes. Subsequently, the surviving h-iPSCs were sorted to isolate monoclonal populations using a fluorescence-activated cell sorter (FACS) in a 96-well plate format. After an additional two weeks of culture, only the h-iPSC colonies demonstrating ETV2-P2A-EYFP expression in response to 1000 ng/mL doxycycline were preserved for future experimental use.

#### Cell culture

h-iPSCs were cultured and passaged without antibiotics in mTeSR Plus medium (STEMCELL Technologies, 100-0276) on 6-well tissue-culture plates coated with hESC-Qualified Matrigel (Corning, 354277). For passaging, after reaching 70% confluency, h-iPSCs were washed with phosphate buffered saline (PBS) (Gibco, 14190250) and dissociated using ReLeSR (STEMCELL Technologies, 100-0483) following the product instruction. h-iPSCs were passaged as small cell aggregates in 1:8 ratio supplemented with 10 µM ROCK inhibitor Y-27632 (STEMCELL Technologies, 72304) for 1 day, followed by complete media change of mTeSR Plus medium every other day. For cryo-preservation, cells were also dissociated with ReLeSR, and resuspended in CryoStor CS10 (STEMCELL Technologies, 07957) as small cell aggregates in 1:8 ratio. Vials were frozen using a CoolCell LX Freezing Container in −80 °C overnight, and subsequently stored in liquid nitrogen for long-term storage.

Human umbilical vein endothelial cells (HUVECs) and normal human lung fibroblasts (HLFs) were purchased from Lonza and cultured in VascuLife VEGF Endothelial Medium (Lifeline Cell Technology) and FibroLife S2 Fibroblast Medium (Lifeline Cell Technology), respectively. HUVECs were transduced to express cytoplasmic GFP, as described earlier (63). Pericytes (PCs) and astrocytes (ACs) were purchased from ScienCell, cultured in corresponding growth medium (ScienCell) on a poly-l-lysine (Sigma) coated flask. All the cells were maintained in a humidified incubator (37 °C, 5% CO_2_), with the culture medium replenished every 2 days. All cell types were used between passages 6–8.

Skov3 tumor cells expressing red fluorescent protein (RFP) were cultured in McCoy’s 5A (Modified) Medium with 10% FBS (Thermo Fisher). Peripheral blood mononuclear cells (PBMCs) and monocytes were isolated from healthy donor’s blood by the monocyte core at MIT. To generate CAR-T cells, PBMCs were treated with Dynabeads™ Human T-Activator CD3/CD28 for T Cell Expansion and Activation (Thermo Fisher), along with IL-2 (18.3 ng/mL), IL-7 (5 ng/mL), and IL-15 (5 ng/mL) in RPMI1640 with 10% heat inactivated FBS for 6 days.

On day 2, expanded cells were infected with lentivirus of mouse single chain anti-human mesothelin specific chimeric antigen receptor overnight. Dynabeads were removed on day 6, and transfected cells were further cultured for another 2 days without cytokines. On day 9, 99% of cells were CD3 positive (BioLegend) and ready for experiments. Control T cells or CAR-T cells were labeled with CellTracker green (Thermo Scientific) prior to the perfusion experiments.

### Differentiation of h-iPSCs into h-iECs

#### Conventional two step method

We followed the protocol established by Orlova et with slight modifications (12). h-iPSCs were dissociated into small cell aggregates using ReLeSR and plated on Matrigel coated 6-well plate with 1:60 ratio in mTeSR Plus medium supplemented with 10 µM Y-27632. The next day, culture medium was changed to mTeSR Plus without Y-27632. 48 h later (day 0), mTeSR Plus was replaced by mesoderm induction medium consisting of VEGF-A (50 ng ml^−1^), FGF-2 (10 ng ml^−1^), BMP4 (30 ng ml^−1^) and CHIR (4 mM) in STEMdiff APEL2 medium (STEMCELL Technologies, 05275). After 2 days, medium was changed to vascular specification medium consisting of VEGF-A (50 ng ml^−1^) and SB431542 (20 mM) in APEL2 medium. vascular specification medium was replenished at days 6 and 9.

#### Two step method with ETV2 activation

Protocols were modified according to Wang et al (25). h-iPSCs were dissociated into single cells with Accutase and plated on Matrigel coated 6-well plate with 250,000 cells per well in mTeSR Plus medium supplemented with 10 µM ROCK inhibitor Y-27632. 24 h later (day 1), the medium was changed to mesoderm induction medium consisting of basal medium supplemented with 6 µM CHIR99021 (Sigma-Aldrich, SML1046). Basal medium was prepared by adding 1× GlutaMax supplement (Thermo Fisher Scientific, 35050061) and L-Ascorbic acid (60 µg/ml, Sigma-Aldrich, A8960) into Advanced Dulbecco’s modified Eagle’s medium (DMEM)/ F12 (Thermo Fisher Scientific,12634010). Mesoderm induction medium was replenished again on day 2. On day 3, the medium was changed to vascular specification medium supplemented with 3 μg ml^−1^ doxycycline. Vascular specification medium consisting of basal medium supplemented with VEGF-A (50 ng/ml, PeproTech, 100-20), fibroblast growth factor 2 (FGF-2, 50 ng/ml, PeproTech, 100-18B), EGF (10 ng/ml, PeproTech, AF-100-15), and 10 µM SB431542 (Selleckchem, S1067). On day 4, vascular specification medium was replenished without doxycycline. On day 5, h-iECs were dissociated into single cells with Accutase and plated on 1% gelatin coated T-175 flask in expansion medium, which is consisting of VascuLife VEGF Endothelial Medium supplemented with 8% Tetracycline Free FBS (Takara Bio USA, 631105) and 10 µM SB431542. Expansion medium was replenished every other day.

#### Flow cytometry

The expressions of CD31, CD34, VEGFR2, and UEA-I were tested by Flow cytometry (BD LSR-II). Cells were stained with PE anti-human CD31 antibody (Biolegend, 1:100), Ulex Europaeus Agglutinin I DyLight 649 (Vector Laboratories, 1:200), APC anti-human CD34 antibody (Biolegend, 1:100), or PE anti-human CD309 antibody (Biolegend, 1:100) on ice for 15 min. Cells were then washed twice with MACS buffer and were ready to be examined under flow cytometry. DAPI was used to exclude dead cells.

#### Purification of h-iECs

For h-iECs differentiated with the conventional 2-step method. On day 10, h-iECs were dissociated into single cells and sorted via fluorescence activated cell sorting (FACS) using CD31 and CD34 signals. The purified h-iECs were then expanded on 1% gelatin coated T75 flask, cultured in expansion medium, which is consisting of VascuLife VEGF Endothelial Medium supplemented with 8% FBS and 10 µM SB431542. For h-iECs differentiated with 2-step method with ETV2 activation, the differentiation efficiency is very high (>95%), so the differentiated cells were not purified. Instead, those cells were used directly to form MVNs after expansion.

#### RNA-seq analysis

Samples from 2D cultures of three parental h-iPSC lines, the corresponding h-iECs generated using our protocol, as well as primary HUVECs and hBMVECs, were analyzed. The groups of PGP1 and iPS01 iPSC lines consist of two biological replicates, while all other groups consist of three biological replicates. Total RNA extraction, library preparation, and sequencing were performed at the Integrated Genomics and Bioinformatics Core in the Koch Institute for integrative Cancer Research at MIT (MA, USA). Total RNA was extracted using TRIzol reagent (Thermo Fisher Scientific) and then cleaned using Perkin Elmer Chemagic360. Sequencing libraries were prepared with the NEBNext Ultra II Directional RNA Library Prep Kit for Illumina (E7760; New England Biolabs). Sequencing was performed using a 100M paired-end configuration on the Singular G4 platform.

Raw data were processed with TrimGalore (version 0.6.10) to remove adapters and low-quality sequences, followed by quality control using FastQC (version 1.24.1) and MultiQC (version 1.24.1). Reads were aligned and quantified using HISAT2 (version 2.2.1), Samtools (version 1.20), and StringTie (version 1.20) against the human genome hg38. Differential expression analysis was performed using DESeq2 (version 1.44.0) on counts, with thresholds at fold change >2 and q-value <0.05. Hierarchical clustering and principal component analysis (PCA) were conducted on rlog variance–stabilized reads (R version 4.4.1). Correlation heatmaps and differential gene heatmaps were produced with the pheatmap package (version 1.0.12). RNA-seq data used in this study are deposited in the BioProject database with BioProject ID PRJNA1176017.

#### Device design and fabrication

Two types of microfluidic devices were used for this study: (i) A commercially available microfluidic 3-channel chip, with one gel channel and two media channels (idenTx 3, AIM Biotech). (ii) A three-channel device, featuring a circular central gel channel with 5 mm diameter and two adjacent medium channels, enabling formation of MVNs spanning over a much larger area (Figure 3b). A partial wall was designed to separate the central gel channel and side medium channels, allowing for surface-tension-assisted filling of cell-laden fibrin gels. Molds of the devices were designed in AutoCAD (Autodesk, Inc), and imported in Fusion 360 (Autodesk, Inc) to generate corresponding tool paths, following by milling delrin block with a micro-CNC milling machine (Bantam Tools). Polydimethylsiloxane (PDMS) based microfluidic devices were then fabricated using the procedure outlined in our previous protocols with the molds.

#### Microvascular Network Formation

Cells were cultured to near-confluence prior to detachment, and seeded into the chip as previously described. Briefly, cells were concentrated in VascuLife containing thrombin (4 U mL^−1^, Sigma). Cell mixture solution was then further mixed with fibrinogen (3 mg mL^−1^ final concentration, Sigma). The cell-laden fibrin solution was quickly injected into the device through the gel loading port. Various combinations of cells were used in this study to engineer different types of MVNs. For MVNs formed with iPSC-EC/HUVEC and HLFs, the final concentration is 8 M iPSC-EC/HUVEC mL^−1^ and 1 M HLFs mL^−1^. For brain MVNs, the final concentration is 7 M iPSC-EC mL^−1^, 1 M ACs mL^−1^, and 0.5 M PCs mL^−1^. After seeding, the devices were placed in incubator for 15 min to allow complete polymerization, before adding VascuLife as culture media. Culture medium was replenished every 24 h. All devices were kept in incubator with daily change of culture media for 5–7 days, until perfusable MVNs were formed.

#### Microvascular Network Perfusion, Imaging, and Analysis

To confirm the perfusability of MVNs, 40 kDa Texas red dextran solution was perfused into the MVNs by generating a slight pressure gradient across the gel region. Confocal images were then acquired using an Olympus FLUOVIEW FV1200 confocal laser scanning microscope with a 10× objective. Z-stack images were acquired with a 5 µm step size. Morphological parameters were analyzed and quantified based on collapsed z-stack images using AutoTube (64).

#### Measurement of Vascular Permeability

Vascular permeability was measured following published protocols (29, 65). Briefly, 40 kDa Texas red dextran (Invitrogen) was perfused into the microvascular networks by generating a slight pressure gradient across the gel of the device. Confocal images were captured with a 5 µm step size at 0 and 6 min, from which permeability was calculated. Several regions of interest (ROIs) were captured via timelapse volumetric imaging for each device.

#### PBMCs Perfusion and Extravasation

PBMCs were revived with RPMI1640 supplemented with 10% FBS, followed by staining with Cell Tracker Green. After extensive washing, PBMCs were resuspended at the concentration of 1×10^6^ cells/mL in a mixed media composed of 50% RPMI1640 (10% FBS) and 50% Vasculife complete medium, supplemented with or without phytohaemagglutinin (PHA, 10 µg/mL). Subsequently, PBMCs were perfused into MVNs that were stained with DyLight 649 labeled UEA-I. Confocal images were taken 24 hours post perfusion to examine cell extravasation.

#### Formation of Vascularized Tumor Spheroid Model

Skov3 tumor spheroid was formed following our published protocol (36). In short, Skov3 cells were first form a core spheroid (500 cells per spheroid) in a 96-well ultra-low attachment plate (Wako Chemicals, USA). Human lung fibroblasts (500 cells per spheroid) were loaded to the pre-formed tumor spheroid to generate a sequential tumor spheroid. Skov3 tumor spheroids were further cultured in McCoy’s 5A (Modified) Medium supplemented with 10% FBS. MVN seeding followed our published protocol (37). HUVEC (outer layer) solution was pipetted into the gel inlet at a final concentration of 12×10^6^/mL, immediately followed by aspirating from the gel outlet. Another solution with final concentrations of 6×10^6^/mL iPSC-ECs, 1.5×10^6^/mL lung FBs, and one Skvo3 tumor spheroid, was then pipetted into the same chip through the gel outlet. After the hydrogel was polymerized, devices were cultured in Vasculife media. After 7 days, devices were stained with DyLight 649 labeled Ulex Europaeus Agglutinin I (UEA-I, Vector laboratories) to monitor vasculature morphology by confocal microscopy. After imaging, devices were then perfused with medium containing dextran (40 kDa, Thermo Scientific) to test the perfusability of MVNs under confocal microscope.

#### T Cell Perfusion in the Vascularized Tumor Device

Control T cells or CAR-T cells were labeled with CellTracker green (Thermo Scientific) and then perfused into MVNs at 10^6^/mL cell density in 20 μL RPMI1640 per device. 30 mins post perfusion, culture media made of 50% Vasculife and 50% RPMI1640 were added. Half of the culture media in the device was replaced with fresh media daily over 3 days. On day 11, conditioned media were collected for interferon gamma (IFNγ) detection (R&D systems). Devices were then stained with DAPI to detect dead cells in vascularized tumor spheroids. After washing, devices were imaged directly using confocal microscopy.

#### Vascularized Liver Organoid Culture

PGP1-GATA6 and Alstem iPS01-ETV2 were dissociated when h-iPSC monolayers reach 70– 80% confluency with Accutase (Sigma-Aldrich, SCR005). Cells were then resuspended in mTeSR Plus medium and centrifuged at 200 g for 5 min. Total of 1.2 × 10^4^ cells were seeded into each well of 96-well Round Bottom Ultra-Low Attachment Microplate (Corning, 7007) to form embryoid bodies (EB), with 150 µL mTeSR Plus supplemented with 10 µM Y-27632 and 1 μg ml^−1^ doxycycline. Three groups with GATA6: ETV2 ratio of 10:0, 8:2, 6:4 were used in this study. The day after aggregation, the culture medium was replaced with fresh mTeSR Plus without Y-27632. mTeSR Plus with doxycycline was used for the first 5 days and replenished daily. On day 6, culture media was changed to 1 to 1 mixed VascuLife and APEL2 medium, without doxycycline, and replenished every other day. Organoids were cultured until day 22. As control experiment, PGP1-GATA6 and HUVECs were mixed in 8:2 ratio and cultured under the same conditions.

#### Immunofluorescence Staining

For immunofluorescent staining, 2D cell culture and various MVNs were fixed by 4% paraformaldehyde. After thorough washing with PBS, cells were permeabilized with 0.2% Triton X-100 and then blocked with 5% Donkey serum (Millipore Sigma, D9663). Subsequently, cells or MVNs were stained with primary antibody on a rocker at 4 °C overnight, following by extensive wash with PBS on a rocker at room temperature, the cells or MVNs were then stained with secondary antibody on a rocker at 4 °C overnight. Organoids were fixed by 4% paraformaldehyde overnight. After thorough washing with PBS, organoids were permeabilized with 2% Triton X-100 and then blocked with 10% Donkey serum for 1 day. Subsequently, organoids were stained with primary antibody on a rocker at 4 °C for 2 days with antibody dilution buffer (1% donkey serum, 0.2% Triton-X 100 in PBS), following by extensive wash with PBS on a rocker at room temperature. The organoids were then incubated with secondary antibody on a rocker at 4°C for 1 day. Primary antibodies used in this study are : Brachyury (R&D Systems, AF2085, 1:100, 5 µg mL^−1^), ETV2 (abcam, ab181847 1:500), CD31 (abcam, ab32457, 1:200), ZO-1 (Thermofisher, 61-7300, 1:100), VE-Cadherin (Biolegend, 348501, 1:100), Collagen IV (Thermofisher, MA1-22148, 1:500), S-100b (Sigma, S2532, 1:200), PDGFR (abcam, ab32570, 1:200), CEBPa (R&D systems, AF7094, 2 µg mL^−1^), Desmin (Thermofisher, MA5-13259, 1:100). Secondary antibodies used in this study are: Donkey anti-Mouse Alexa Fluo 488, Donkey anti-Rabbit Alexa Fluo 568, Donkey anti-Goat Alexa Fluo 568, Donkey anti-Sheep Alexa Fluo 647, Donkey anti-Mouse Alexa Fluo 647 (Invitrogen, 1:200). Ulex Europaeus Agglutinin I (UEA I) was also used in this study for MVNs live stain, before perfusing PBMCs or T cells.

#### Vascularized Liver Organoid tissue clearing and imaging

The immunostained organoids were cleared with RapiClear 1.49 (SUNJin Lab, RC149001) at room temperature overnight. The next day, cleared organoids were mounted with fresh RapiClear reagent in iSpacer (SUNJin Lab, IS008) microchambers, with thickness of 0.5 mm. The chambers were sealed by gently pressing coverslip around the iSpacer. Confocal images were then acquired using an Olympus FLUOVIEW FV1200 confocal laser scanning microscope with a 10× objective. Z-stack images were acquired with a 5 µm step size.

#### Morphological Analysis of Vascularized Liver Organoid

3D volume of liver organoids was calculated based on DAPI signals from 3D confocal z-stack using customized Matlab code. Morphological parameters of vascular network within organoids were calculated from CD31 signals in confocal z-stack with ImageJ. Briefly, the image stack was thresholded, and the 3D skeleton was subsequently extracted using skeletonize (2D/3D) plugin. The 3D skeleton was further analyzed to get the total number of endpoints (having only one neighboring branch connected to them) and junctions (having more than two neighboring branches connected to them), the total vessel path length (total path length of all identified skeleton), and the dominant vessel path length (the longest skeleton path). We further define dominant skeleton ratio (proportion of the longest skeleton path relative to the total path length of all skeletons) and branching factor (the ratio between junction numbers and endpoint numbers) to better characterize the connectivity of vascular network within organoids.

#### Measurement of Urea Production

QuantiChrom Urea Assay Kit (BioAssay Systems, DIUR-100) was used to measure urea concentration following product instructions. Briefly, culture medium from day 22 organoid were sampled and mixed with the working reagent. After incubating 50 mins at room temperature, optical densities (OD) at 430 nm were read using BioTek Synergy Neo2 Reader. The urea concentrations were then calculated based on OD of the sample, the blank sample, and standard samples.

#### Statistics

All bar plots are shown as mean ± SD and plotted with Prism (GraphPad). All data representation details are provided in corresponding figure captions. Statistical significance was assessed using t-test performed in Matlab (MathWorks). * denotes p<0.05, *** denotes p < 0.01.

## Supporting Information

Supporting Information is available from the Wiley Online Library or from the author.

## Acknowledgements

This work was supported by the Strategic Priority Research Program of the Chinese Academy of Sciences (XDB0820000) and Wellcome Leap HOPE Program. Shun Zhang and Zhengpeng Wan contributed equally to this work. PBMCs were kindly provided by Prof. Bryan Bryson. Francesca M. Pramotton was supported by the Postdoc. Mobility fellowship (P500PT 211085).

## References

1. Zhang B, Korolj A, Lai BFL, Radisic M. Advances in organ-on-a-chip engineering. Nature Reviews Materials. 2018;3(8):257–78.

2. Rossi G, Manfrin A, Lutolf MP. Progress and potential in organoid research. Nat Rev Genet. 2018;19(11):671–87.

3. Lancaster MA, Knoblich JA. Organogenesis in a dish: modeling development and disease using organoid technologies. Science. 2014;345(6194):1247125.

4. Hofer M, Lutolf MP. Engineering organoids. Nat Rev Mater. 2021;6(5):402–20.

5. Leung CM, de Haan P, Ronaldson-Bouchard K, Kim G-A, Ko J, Rho HS, et al. A guide to the organ-on-a-chip. Nature Reviews Methods Primers. 2022;2(1):33.

6. Bhatia SN, Ingber DE. Microfluidic organs-on-chips. Nat Biotechnol. 2014;32(8):760–72.

7. Zhang S, Wan Z, Kamm RD. Vascularized organoids on a chip: strategies for engineering organoids with functional vasculature. Lab Chip. 2021;21(3):473–88.

8. Grebenyuk S, Ranga A. Engineering Organoid Vascularization. Front Bioeng Biotechnol. 2019;7:39.

9. Zhang S, Kan EL, Kamm RD. Integrating functional vasculature into organoid culture: A biomechanical perspective. APL Bioeng. 2022;6(3):030401.

10. Samuel R, Duda DG, Fukumura D, Jain RK. Vascular diseases await translation of blood vessels engineered from stem cells. Sci Transl Med. 2015;7(309):309rv6.

11. Patsch C, Challet-Meylan L, Thoma EC, Urich E, Heckel T, O’Sullivan JF, et al. Generation of vascular endothelial and smooth muscle cells from human pluripotent stem cells. Nat Cell Biol. 2015;17(8):994–1003.

12. Orlova VV, van den Hil FE, Petrus-Reurer S, Drabsch Y, Ten Dijke P, Mummery CL. Generation, expansion and functional analysis of endothelial cells and pericytes derived from human pluripotent stem cells. Nat Protoc. 2014;9(6):1514–31.

13. Paik DT, Tian L, Lee J, Sayed N, Chen IY, Rhee S, et al. Large-Scale Single-Cell RNA-Seq Reveals Molecular Signatures of Heterogeneous Populations of Human Induced Pluripotent Stem Cell-Derived Endothelial Cells. Circ Res. 2018;123(4):443–50.

14. Hu S, Zhao MT, Jahanbani F, Shao NY, Lee WH, Chen H, et al. Effects of cellular origin on differentiation of human induced pluripotent stem cell-derived endothelial cells. JCI Insight. 2016;1(8).

15. Kataoka H, Hayashi M, Nakagawa R, Tanaka Y, Izumi N, Nishikawa S, et al. Etv2/ER71 induces vascular mesoderm from Flk1+PDGFRalpha+ primitive mesoderm. Blood. 2011;118(26):6975–86.

16. Oh SY, Kim JY, Park C. The ETS Factor, ETV2: a Master Regulator for Vascular Endothelial Cell Development. Mol Cells. 2015;38(12):1029–36.

17. Ng AHM, Khoshakhlagh P, Rojo Arias JE, Pasquini G, Wang K, Swiersy A, et al. A comprehensive library of human transcription factors for cell fate engineering. Nat Biotechnol. 2021;39(4):510–9.

18. Gong W, Das S, Sierra-Pagan JE, Skie E, Dsouza N, Larson TA, et al. ETV2 functions as a pioneer factor to regulate and reprogram the endothelial lineage. Nat Cell Biol. 2022;24(5):672–84.

19. Morita R, Suzuki M, Kasahara H, Shimizu N, Shichita T, Sekiya T, et al. ETS transcription factor ETV2 directly converts human fibroblasts into functional endothelial cells. Proc Natl Acad Sci U S A. 2015;112(1):160–5.

20. Lange L, Hoffmann D, Schwarzer A, Ha TC, Philipp F, Lenz D, et al. Inducible Forward Programming of Human Pluripotent Stem Cells to Hemato-endothelial Progenitor Cells with Hematopoietic Progenitor Potential. Stem Cell Reports. 2020;14(1):122–37.

21. Linville RM, Sklar MB, Grifno GN, Nerenberg RF, Zhou J, Ye R, et al. Three-dimensional microenvironment regulates gene expression, function, and tight junction dynamics of iPSC-derived blood-brain barrier microvessels. Fluids Barriers CNS. 2022;19(1):87.

22. Lu TM, Houghton S, Magdeldin T, Duran JGB, Minotti AP, Snead A, et al. Pluripotent stem cell-derived epithelium misidentified as brain microvascular endothelium requires ETS factors to acquire vascular fate. Proc Natl Acad Sci U S A. 2021;118(8).

23. Zhang H, Yamaguchi T, Kokubu Y, Kawabata K. Transient ETV2 Expression Promotes the Generation of Mature Endothelial Cells from Human Pluripotent Stem Cells. Biol Pharm Bull. 2022;45(4):483–90.

24. Skylar-Scott MA, Huang JY, Lu A, Ng AHM, Duenki T, Liu S, et al. Orthogonally induced differentiation of stem cells for the programmatic patterning of vascularized organoids and bioprinted tissues. Nat Biomed Eng. 2022;6(4):449–62.

25. Wang K, Lin RZ, Hong X, Ng AH, Lee CN, Neumeyer J, et al. Robust differentiation of human pluripotent stem cells into endothelial cells via temporal modulation of ETV2 with modified mRNA. Sci Adv. 2020;6(30):eaba7606.

26. Luo AC, Wang J, Wang K, Zhu Y, Gong L, Lee U, et al. A streamlined method to generate endothelial cells from human pluripotent stem cells via transient doxycycline-inducible ETV2 activation. Angiogenesis. 2024.

27. Palikuqi B, Nguyen DT, Li G, Schreiner R, Pellegata AF, Liu Y, et al. Adaptable haemodynamic endothelial cells for organogenesis and tumorigenesis. Nature. 2020;585(7825):426–32.

28. Guye P, Ebrahimkhani MR, Kipniss N, Velazquez JJ, Schoenfeld E, Kiani S, et al. Genetically engineering self-organization of human pluripotent stem cells into a liver bud-like tissue using Gata6. Nat Commun. 2016;7:10243.

29. Hajal C, Offeddu GS, Shin Y, Zhang S, Morozova O, Hickman D, et al. Engineered human blood-brain barrier microfluidic model for vascular permeability analyses. Nat Protoc. 2022;17(1):95–128.

30. Winkelman MA, Kim DY, Kakarla S, Grath A, Silvia N, Dai G. Interstitial flow enhances the formation, connectivity, and function of 3D brain microvascular networks generated within a microfluidic device. Lab Chip. 2021;22(1):170–92.

31. Campisi M, Shin Y, Osaki T, Hajal C, Chiono V, Kamm RD. 3D self-organized microvascular model of the human blood-brain barrier with endothelial cells, pericytes and astrocytes. Biomaterials. 2018;180:117–29.

32. Haase K, Offeddu GS, Gillrie MR, Kamm RD. Endothelial Regulation of Drug Transport in a 3D Vascularized Tumor Model. Adv Funct Mater. 2020;30(48).

33. Hachey SJ, Movsesyan S, Nguyen QH, Burton-Sojo G, Tankazyan A, Wu J, et al. An in vitro vascularized micro-tumor model of human colorectal cancer recapitulates in vivo responses to standard-of-care therapy. Lab Chip. 2021;21(7):1333–51.

34. Nguyen HT, Kan EL, Humayun M, Gurvich N, Offeddu GS, Wan Z, et al. Patient-specific vascularized tumor model: Blocking monocyte recruitment with multispecific antibodies targeting CCR2 and CSF-1R. Biomaterials. 2025;312:122731.

35. Offeddu GS, Cambria E, Shelton SE, Haase K, Wan Z, Possenti L, et al. Personalized Vascularized Models of Breast Cancer Desmoplasia Reveal Biomechanical Determinants of Drug Delivery to the Tumor. Adv Sci (Weinh). 2024;11(38):e2402757.

36. Wan Z, Floryan MA, Coughlin MF, Zhang S, Zhong AX, Shelton SE, et al. New Strategy for Promoting Vascularization in Tumor Spheroids in a Microfluidic Assay. Adv Healthc Mater. 2023;12(14):e2201784.

37. Wan Z, Zhong AX, Zhang S, Pavlou G, Coughlin MF, Shelton SE, et al. A Robust Method for Perfusable Microvascular Network Formation In Vitro. Small Methods. 2022;6(6):e2200143.

38. Cakir B, Xiang Y, Tanaka Y, Kural MH, Parent M, Kang YJ, et al. Engineering of human brain organoids with a functional vascular-like system. Nat Methods. 2019;16(11):1169–75.

39. Takebe T, Sekine K, Enomura M, Koike H, Kimura M, Ogaeri T, et al. Vascularized and functional human liver from an iPSC-derived organ bud transplant. Nature. 2013;499(7459):481–4.

40. Takahashi Y, Sekine K, Kin T, Takebe T, Taniguchi H. Self-Condensation Culture Enables Vascularization of Tissue Fragments for Efficient Therapeutic Transplantation. Cell Rep. 2018;23(6):1620–9.

41. Pham MT, Pollock KM, Rose MD, Cary WA, Stewart HR, Zhou P, et al. Generation of human vascularized brain organoids. Neuroreport. 2018;29(7):588–93.

42. Lin Y, Gil CH, Yoder MC. Differentiation, Evaluation, and Application of Human Induced Pluripotent Stem Cell-Derived Endothelial Cells. Arterioscler Thromb Vasc Biol. 2017;37(11):2014–25.

43. Ewald ML, Chen YH, Lee AP, Hughes CCW. The vascular niche in next generation microphysiological systems. Lab Chip. 2021;21(17):3244–62.

44. Harding A, Cortez-Toledo E, Magner NL, Beegle JR, Coleal-Bergum DP, Hao D, et al. Highly Efficient Differentiation of Endothelial Cells from Pluripotent Stem Cells Requires the MAPK and the PI3K Pathways. Stem Cells. 2017;35(4):909–19.

45. Sahara M, Hansson EM, Wernet O, Lui KO, Spater D, Chien KR. Manipulation of a VEGF-Notch signaling circuit drives formation of functional vascular endothelial progenitors from human pluripotent stem cells. Cell Res. 2014;24(7):820–41.

46. Geudens I, Gerhardt H. Coordinating cell behaviour during blood vessel formation. Development. 2011;138(21):4569–83.

47. Bergers G, Song S. The role of pericytes in blood-vessel formation and maintenance. Neuro Oncol. 2005;7(4):452–64.

48. Semenza GL. Vasculogenesis, angiogenesis, and arteriogenesis: mechanisms of blood vessel formation and remodeling. J Cell Biochem. 2007;102(4):840–7.

49. Bonanini F, Kurek D, Previdi S, Nicolas A, Hendriks D, de Ruiter S, et al. In vitro grafting of hepatic spheroids and organoids on a microfluidic vascular bed. Angiogenesis. 2022;25(4):455–70.

50. Song L, Yuan X, Jones Z, Griffin K, Zhou Y, Ma T, et al. Assembly of Human Stem Cell-Derived Cortical Spheroids and Vascular Spheroids to Model 3-D Brain-like Tissues. Sci Rep. 2019;9(1):5977.

51. Wang L, Xu W, Zhang S, Gundberg GC, Zheng CR, Wan Z, et al. Sensing and guiding cell-state transitions by using genetically encoded endoribonuclease-mediated microRNA sensors. Nat Biomed Eng. 2024.

52. Zhang S, Wan Z, Pavlou G, Zhong AX, Xu L, Kamm RD. Interstitial flow promotes the formation of functional microvascular networks in vitro through upregulation of matrix metalloproteinase-2. Adv Funct Mater. 2022;32(43).

53. Galie PA, Nguyen DH, Choi CK, Cohen DM, Janmey PA, Chen CS. Fluid shear stress threshold regulates angiogenic sprouting. Proc Natl Acad Sci U S A. 2014;111(22):7968–73.

54. Kim S, Chung M, Ahn J, Lee S, Jeon NL. Interstitial flow regulates the angiogenic response and phenotype of endothelial cells in a 3D culture model. Lab Chip. 2016;16(21):4189–99.

55. Helm CL, Fleury ME, Zisch AH, Boschetti F, Swartz MA. Synergy between interstitial flow and VEGF directs capillary morphogenesis in vitro through a gradient amplification mechanism. Proc Natl Acad Sci U S A. 2005;102(44):15779–84.

56. Homan KA, Gupta N, Kroll KT, Kolesky DB, Skylar-Scott M, Miyoshi T, et al. Flow-enhanced vascularization and maturation of kidney organoids in vitro. Nat Methods. 2019;16(3):255–62.

57. Elcheva I, Brok-Volchanskaya V, Kumar A, Liu P, Lee JH, Tong L, et al. Direct induction of haematoendothelial programs in human pluripotent stem cells by transcriptional regulators. Nat Commun. 2014;5:4372.

58. Barger CJ, Chee L, Albahrani M, Munoz-Trujillo C, Boghean L, Branick C, et al. Co-regulation and function of FOXM1/RHNO1 bidirectional genes in cancer. Elife. 2021;10.

59. Guye P, Li Y, Wroblewska L, Duportet X, Weiss R. Rapid, modular and reliable construction of complex mammalian gene circuits. Nucleic Acids Res. 2013;41(16):e156.

60. Duportet X, Wroblewska L, Guye P, Li Y, Eyquem J, Rieders J, et al. A platform for rapid prototyping of synthetic gene networks in mammalian cells. Nucleic Acids Res. 2014;42(21):13440–51.

61. Chen F, LoTurco J. A method for stable transgenesis of radial glia lineage in rat neocortex by piggyBac mediated transposition. J Neurosci Methods. 2012;207(2):172–80.

62. Yusa K, Zhou L, Li MA, Bradley A, Craig NL. A hyperactive piggyBac transposase for mammalian applications. Proc Natl Acad Sci U S A. 2011;108(4):1531–6.

63. Wan Z, Zhang S, Zhong AX, Shelton SE, Campisi M, Sundararaman SK, et al. A robust vasculogenic microfluidic model using human immortalized endothelial cells and Thy1 positive fibroblasts. Biomaterials. 2021;276:121032.

64. Montoya-Zegarra JA, Russo E, Runge P, Jadhav M, Willrodt AH, Stoma S, et al. AutoTube: a novel software for the automated morphometric analysis of vascular networks in tissues. Angiogenesis. 2019;22(2):223–36.

65. Offeddu GS, Haase K, Gillrie MR, Li R, Morozova O, Hickman D, et al. An on-chip model of protein paracellular and transcellular permeability in the microcirculation. Biomaterials. 2019;212:115–25.

