## Supplemental data for "ETV2 mediated differentiation of human pluripotent stem cells results in functional endothelial cells for engineering advanced vascularized microphysiological models"

S. Zhang, C. Wu, J. Zhang
State Key Laboratory of Organ Regeneration and Reconstruction, Institute of Zoology, Chinese Academy of Sciences, Beijing 100101, China
Beijing Institute for Stem Cell and Regenerative Medicine, Beijing 100101, China
Z. Wan, L. Wang, X. Wang, M. F. Coughlin, F. Pramotton, R. Weiss, R. D. Kamm
Department of Biological Engineering, Massachusetts Institute of Technology, Cambridge, MA, 02139, USA
M. A. Floryan, R. D. Kamm
Department of Mechanical Engineering Massachusetts Institute of Technology Cambridge, MA 02139, USA
L. Wang
Bioengineering Department, Northeastern University, Boston, MA, 02115, USA
L. Xu
Ragon Institute of MGH, MIT and Harvard, Cambridge, MA, 02139, USA
R. Weiss
Synthetic Biology Center, Massachusetts Institute of Technology, Cambridge, MA, 02139, USA
*corresponding authors, [#]contributed equally to this work


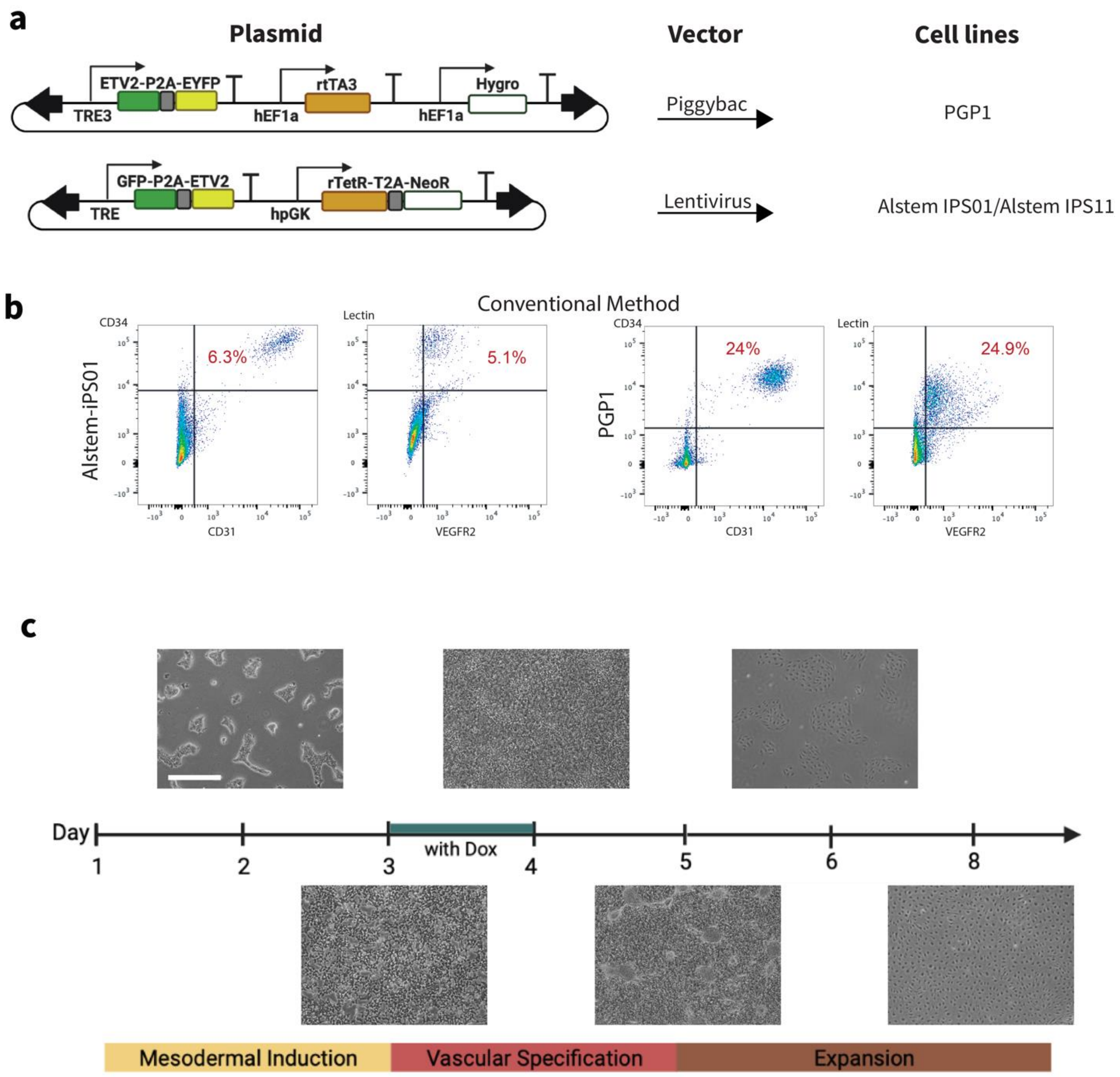


Figure S1. Differentiation of h-iECs from h-iPSCs with engineered inducible ETV2 with optimized protocol. (a) Generation of various h-iPSC lines with inducible ETV2 using Piggybac or lentivirus. (b) Differentiation efficiency of h-iPSCs into $CD31^+/CD34^+/VEGFR2^+/$ UEA-I $^+$ h-iECs with conventional two step method by flow cytometry. (c) Bright field images showing differentiation process of h-iECs from h-iPSCs with engineered inducible ETV2. Scale bar is 500 μm.

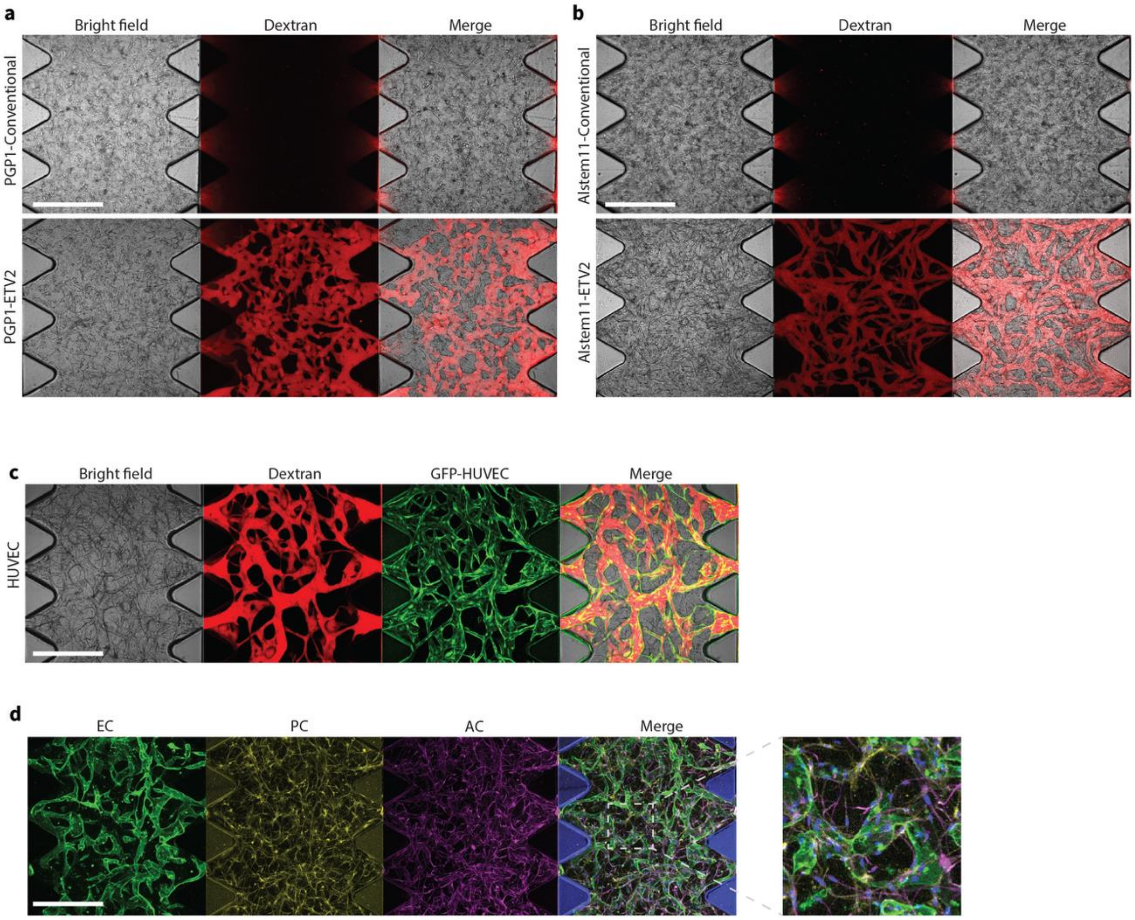
a
Bright field
Dextran
Merge
PGP1-Conventional
PGP1-ETV2
b
Bright field
Dextran
Merge
Alstem11-Conventional
Alstem11-ETV2
c
Bright field
Dextran
GFP-HUVEC
Merge
HUVEC
d
EC
PC
AC
Merge

Figure S2. Formation of functional MVNs with h-iECs differentiated through transient activation of ETV2. (a) Representative images of MVNs made of h-iECs on day 7. h-iECs were differentiated from PGP1 h-iPSCs using either conventional method or optimized protocol with ETV2 activation. Perfusion test was performed with 40 kDa Texas Red dextran (red). (b) Representative images of MVNs made of h-iECs on day 7. h-iECs were differentiated from Alstem iPS11 h-iPSCs using either conventional method or optimized protocol with ETV2 activation. Perfusion test was performed with 40 kDa Texas Red dextran (red). (c) Representative images of MVNs formed using HUVECs and HLFs at day 7. Red: 40 kDa Texas Red dextran, Green: GFP HUVECs. (d) Immunofluorescence staining for brain specific MVNs made of h-iECs, human brain pericytes (PCs), and astrocytes (ACs), with zoomed-in view. Green: CD31 staining for h-iECs, magenta: S-100b staining for ACs, yellow: PDGFR staining for PCs. All scale bars are 500 μm.

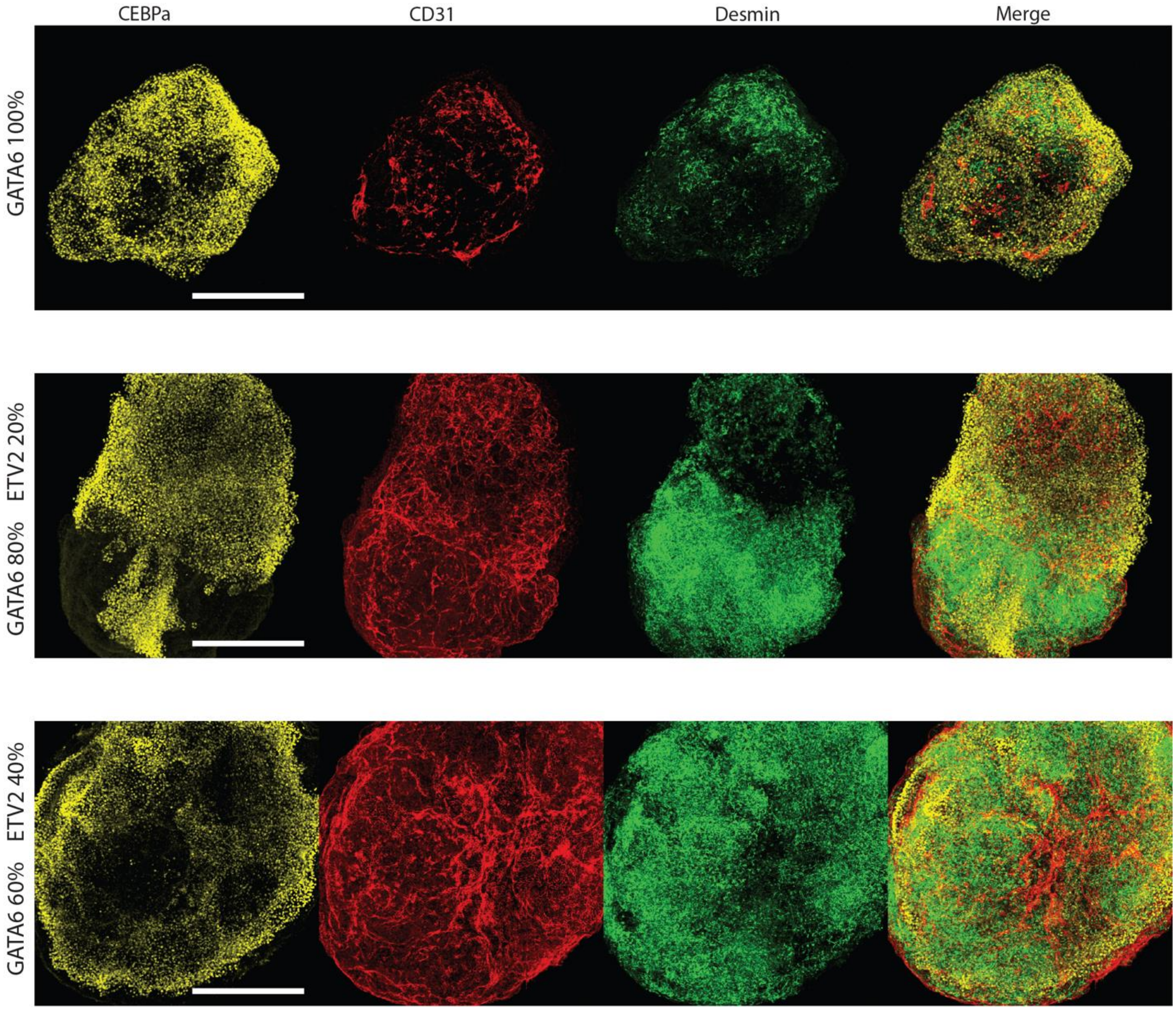


Figure S3. Immunofluorescence staining for liver organoids formed by pooling different ratios of PGP1-GATA6 and PGP1-ETV2. Yellow: CEBPa staining for hepatocytes, Red: CD31 staining for ECs, Green: Desmin staining for stellate cells. Scale bar is 500 μm.

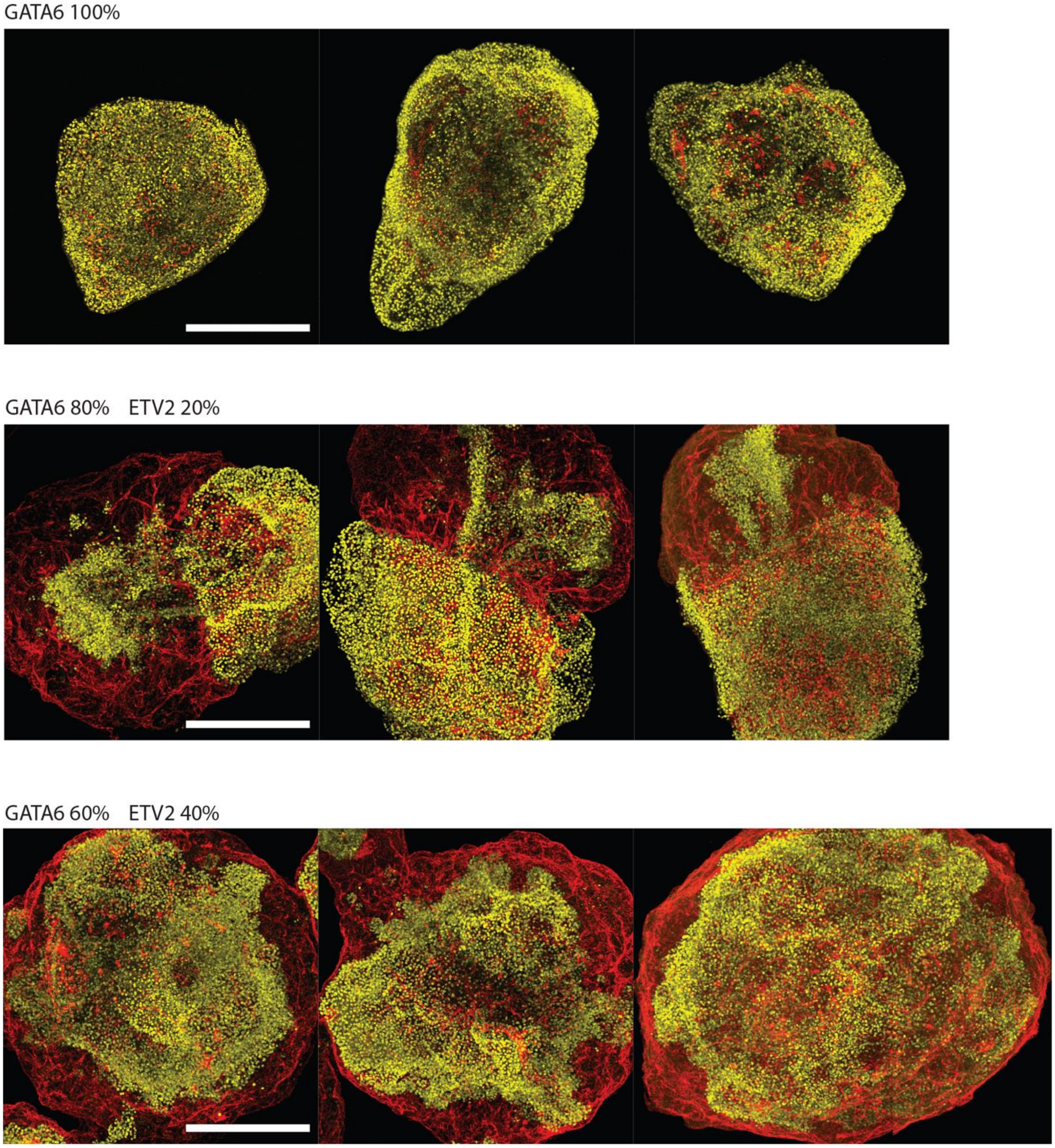


Figure S4. Spatial arrangements of ECs and hepatocytes within liver organoids formed by pooling different ratios of PGP1-GATA6 and PGP1-ETV2. Yellow: CEBPa staining for hepatocytes, Red: CD31 staining for ECs. Scale bars are 500 μm.

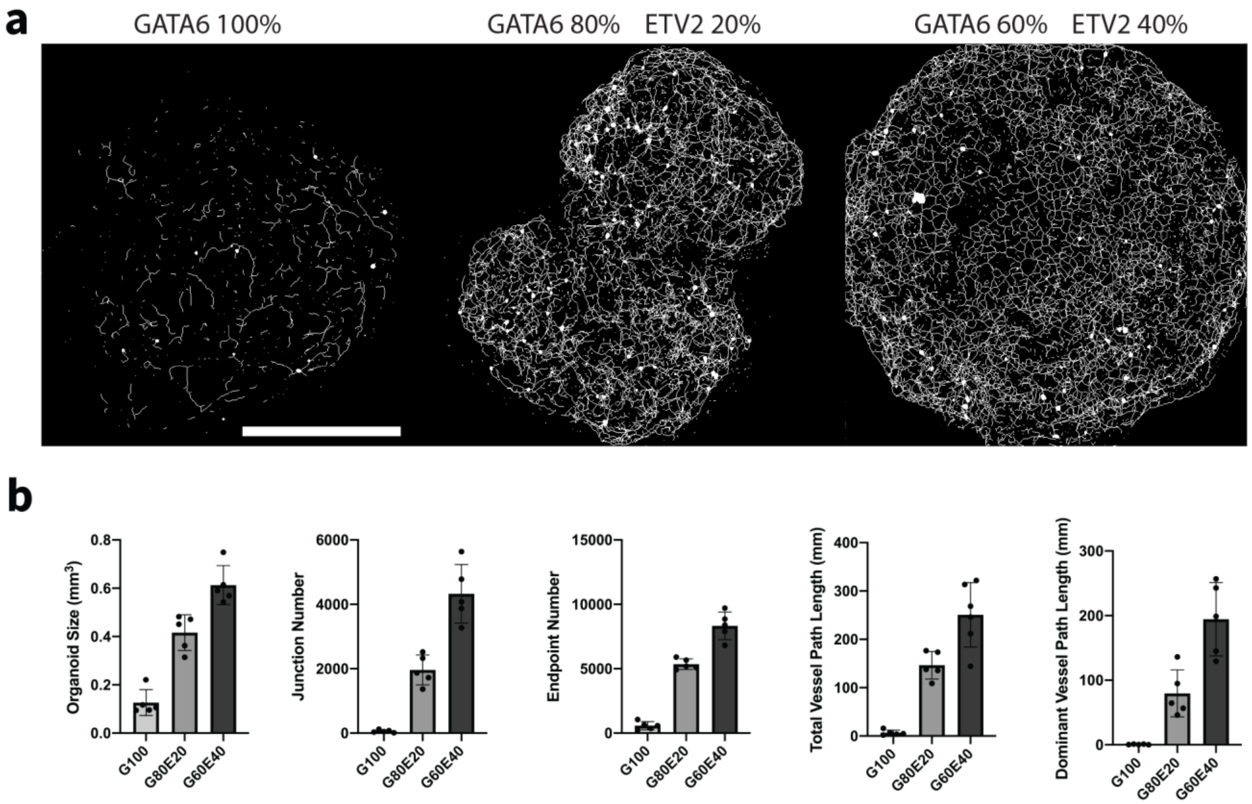


Figure S5. Characterization of vascular networks within liver organoids formed by pooling different ratios of PGP1-GATA6 and PGP1-ETV2. (a) 3D skeletons of vascular networks within liver organoids. Scale bar is 500 μm. (b) Detailed characterization of morphological parameters of vascular networks within liver organoids formed by pooling different ratios of PGP1-GATA6 and PGP1-ETV2.
